# Long-read-based Human Genomic Structural Variation Detection with cuteSV

**DOI:** 10.1101/780700

**Authors:** Tao Jiang, Bo Liu, Yue Jiang, Junyi Li, Yan Gao, Zhe Cui, Yadong Liu, Yadong Wang

## Abstract

Long-read sequencing enables the comprehensive discovery of structural variations (SVs). However, it is still non-trivial to achieve high sensitivity and performance simultaneously due to the complex SV characteristics implied by noisy long reads. Therefore, we propose cuteSV, a sensitive, fast and scalable long-read-based SV detection approach. cuteSV uses tailored methods to collect the signatures of various types of SVs and employs a clustering-and-refinement method to analyze the signatures to implement sensitive SV detection. Benchmarks on real PacBio and ONT datasets demonstrate that cuteSV has better yields and scalability than state-of-the-art tools. cuteSV is available at https://github.com/tjiangHIT/cuteSV.

## Background

Structural variations (SVs) represent genomic rearrangements such as deletions, insertions, inversions, duplications, and translocations whose sizes are larger than 50 bp [1]. As the largest divergences across human genomes [2], SVs are closely related to human diseases (e.g., inherited diseases [3-5] and cancers [6]), evolution (e.g., gene losses and transposon activity [7, 8]), gene regulations (e.g., rearrangements of transcription factors [9]) and other phenotypes (e.g., mating and intrinsic reproductive isolation [10, 11]).

Many efforts have been made to develop short-read-based SV calling approaches [12, 13]. State-of-the-art short-read-based tools use various kinds of methods such as read-depths [14], discordant read-pairs [15], split read alignments [16], local assembly [17] or their combinations [18-20]. They have played important roles in many large-scale genomics studies such as the 1000 Genome Project [1]. However, the relatively low read-length (typically a few hundreds of bps) limits these tools in implementing highly sensitive SV detection [21] and false positives are common in their call sets [22].

With the rapid development of long-read sequencing technologies, such as Pacific Bioscience (PacBio) [23] and Oxford Nanopore Technology (ONT) [24] platforms, read length has greatly increased. The average read length can be over 10 kbp, and long-range spanning information provides the opportunity to comprehensively detect SVs at a high resolution [25]. However, the computational challenge still remains due to the high sequencing error rates of long-reads (typically 5-20%) [26]. Because of the sequence error, the aligners may not be able to produce sensitive, chimeric and heterogeneous alignments for the reads around SV breakpoints, and the aligners can also have highly divergent behaviors for various types of SVs. Therefore, the SV signatures implied by long-read alignments are highly complicated and it is non-trivial to collect and analyze them to implement sensitive detection for various kinds of SVs.

Several long-read-alignment-based SV callers have been proposed in recent years, such as PB-Honey [27], SMRT-SV [28], Sniffles [29], PBSV (https://github.com/PacificBiosciences/pbsv) and SVIM [30]. They use various methods to find evidence of SVs implied by read alignments, such as the identification of local genomic regions with highly divergent alignments, the local assembly and re-alignment of clipped read parts and the clustering of SV-spanning signatures [31]. Moreover, state-of-the-art long-read aligners, such as BLASR [32], NGMLR [29], Minimap2 [33] and PBMM2 (https://github.com/PacificBiosciences/pbmm2), are usually employed for read alignment (as the inputs for the SV callers).

These state-of-the-art long-read-based SV callers have some drawbacks: 1) overall, the sensitivity is not satisfying (i.e., a high sequencing coverage is required and/or some SVs are still hard to detect); 2) some methods such as rMETL [34], rCANID [35] and npInv [36] can only detect a subset or a particular class of SVs due to their specific design; 3) some tools such as PBSV and SMRT-SV are still time-consuming and require plenty of computational resources, and are therefore often not very scalable or suited to many large-scale datasets; 4) some approaches such as SMRT-SV and PB-Honey only support one type of sequencing data (e.g., only for PacBio reads), as they take advantage of the characteristics of the data. These drawbacks currently inhibit the wide use of long-read sequencing data in cutting-edge genomics studies and clinical practices.

Herein, we present cuteSV, a sensitive, fast and scalable long-read-based SV detection approach. This approach has several features: 1) cuteSV has better SVs detection yields than those of state-of-the-art SV callers. Moreover, it has higher sensitivity for low coverage datasets, which is helpful in reducing the cost of sequencing. 2) cuteSV is a versatile SV caller which supports the processing of datasets produced by mainstream platforms with various error rates and the discovery of various types of SVs including deletions, insertions, duplications, inversions and translocations. 3) With its tailored implementation, cuteSV has faster or comparable speed compared to state-of-the-art approaches. 4) cuteSV has outstanding scalability; it can run with low memory use and achieve nearly linear speedup with the number of CPU threads, and is thus very suited to large-scale data analysis tasks. With these features, we believe that cuteSV has a great potential for cutting-edge genomics studies.

## Results

### Overview of cuteSV

Due to the high sequencing error and the complexity of SVs, the SV signals implied by the alignment of SV-spanning long reads are highly complicated, so SV detection approaches must have a strong ability to handle various kinds of cases. Primarily, the following three technical issues should be fully considered.

1. Various types of SVs have different signatures, which can be represented as various combinations of large insertions/deletions and split alignments. SV signature extraction methods have to capture these signals comprehensively. Moreover, due to the scoring systems of read aligners, some large SV events could be divided into several smaller insertions/deletions in a local region. Such signals should be well-handled in order to recover evidence of real SV events.
2. Reads spanning the same SV could have heterogeneous breakpoints in their alignments because of the scoring systems of read aligners. Robust read clustering methods are needed for handling heterogeneous breakpoints in order to cluster reads spanning the same SVs effectively and to prevent multiple false positive clusters for one SV.
3. In some cases, there are multiple SVs in the same local genomics region but belonging to various alleles. It is very difficult to make correct calls for such complex heterozygous SVs. Precise analysis of read alignments is needed to distinguish the signatures of multiple SVs in such loci.

cuteSV is a novel SV detection approach which fulfills the requirements mentioned above. It integrates several specifically designed methods to extract the SV signatures of various kinds of SVs from aligned reads, and it uses a clustering-and-refinement method to thoroughly analyze the collected signatures to call SVs. The approach has three major steps as follows (a schematic illustration is in Figure 1).

**Figure 1.**
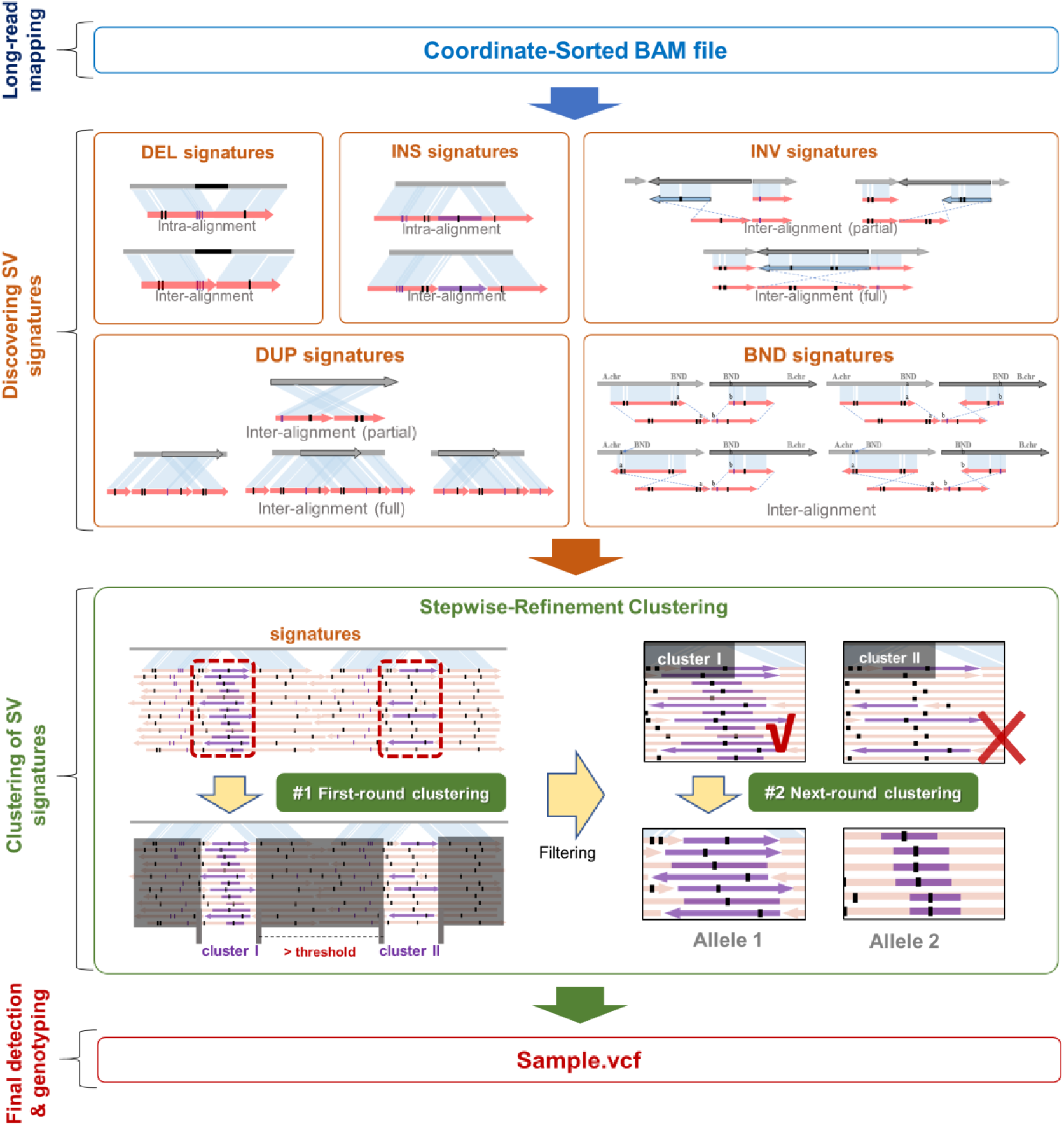
Schematic illustration of the cuteSV approach. The detection of SVs in cuteSV generally comprises 4 steps. In step 1 (long-read mapping), cuteSV supports the sorted-alignment BAM file as input, which is generated from the state-of-the-art long-read mappers. In step 2 (discovering SV signatures), cuteSV collects various type of SV signatures comprehensively from inter- and intra-alignments. In step 3 (clustering of SV signatures), a stepwise-refinement clustering method is employed to sensitively discover accurate SV alleles. In the final step (final detection and genotyping), cuteSV generates the final SV callsets and (optionally) assigns genotypes.

1. cuteSV uses a series of signature extraction methods tailored for various types of SVs. This is one of the two core features of cuteSV that enables the comprehensive collection of SV signatures and the effective recovery of evidence of SVs from complicated and fragile alignments.
2. cuteSV uses a specifically designed clustering-and-refinement approach to cluster the chimerically aligned reads in local regions, and further refines the clusters to precisely distinguish the SV signatures from heterozygous SVs. This is the other core feature of cuteSV that enables it to handle heterogeneous breakpoints in read alignments well and be robust to multi-allelic heterozygous SVs.
3. cuteSV uses several heuristic rules to make SV calls based on clustered SV signatures and to perform high-quality SV genotyping.

Moreover, cuteSV uses a block division-based approach to process input data in a parallel way with multiple CPU threads. Refer to the Materials and methods section for more detailed information about the implementation of cuteSV.

The following sub-sections discuss the benchmark results of cuteSV on real sequencing datasets, and more insights on the features of cuteSV are given in the Discussion section.

### SV detection with HG002 PacBio data

First, we implemented cuteSV and three state-of-the-art long-read-based SV callers (i.e., Sniffles, PBSV and SVIM) on a 69× HG002 PacBio CLR dataset [37] (mean read length: 7938 bp). PBMM2 was employed for read alignment. Moreover, another state-of-the-art aligner, NGMLR, was also employed to investigate the effect of the aligners on SV calling. In this section, the results are based on the alignments produced by PBMM2 unless specifically mentioned otherwise. A high-confidence insertion and deletion callset for this sample made by Genome in a Bottle Consortium (GIAB) [38] was employed as the ground truth. Truvari (https://github.com/spiralgenetics/truvari) was used to assess the precision, recall, and F-measure (F1) of the callsets produced by various approaches.

The yields of the SV callers are shown in Figure 2A. In terms of the 69× CLR data, cuteSV simultaneously achieved the highest precision, recall and F1, all of which were >94% in absolute terms, which is feasible for practical use. The F1 of SVIM (91.10%), PBSV (90.72%) and Sniffles (89.82%) were comparable to each other and slightly lower than that of cuteSV mainly due to their lower recall statistics (i.e., 89.56%, 88.42% and 86.27% for SVIM, PBSV and Sniffles, respectively). For evaluation of genotyping, cuteSV achieved > 90% both on GT-recall and GT-F1. SVIM was the runner-up and reached 85% on all three statistics. PBSV and Sniffles were far behind cuteSV and SVIM due to their relatively poor GT-precision and GT-recall.

**Figure 2.**
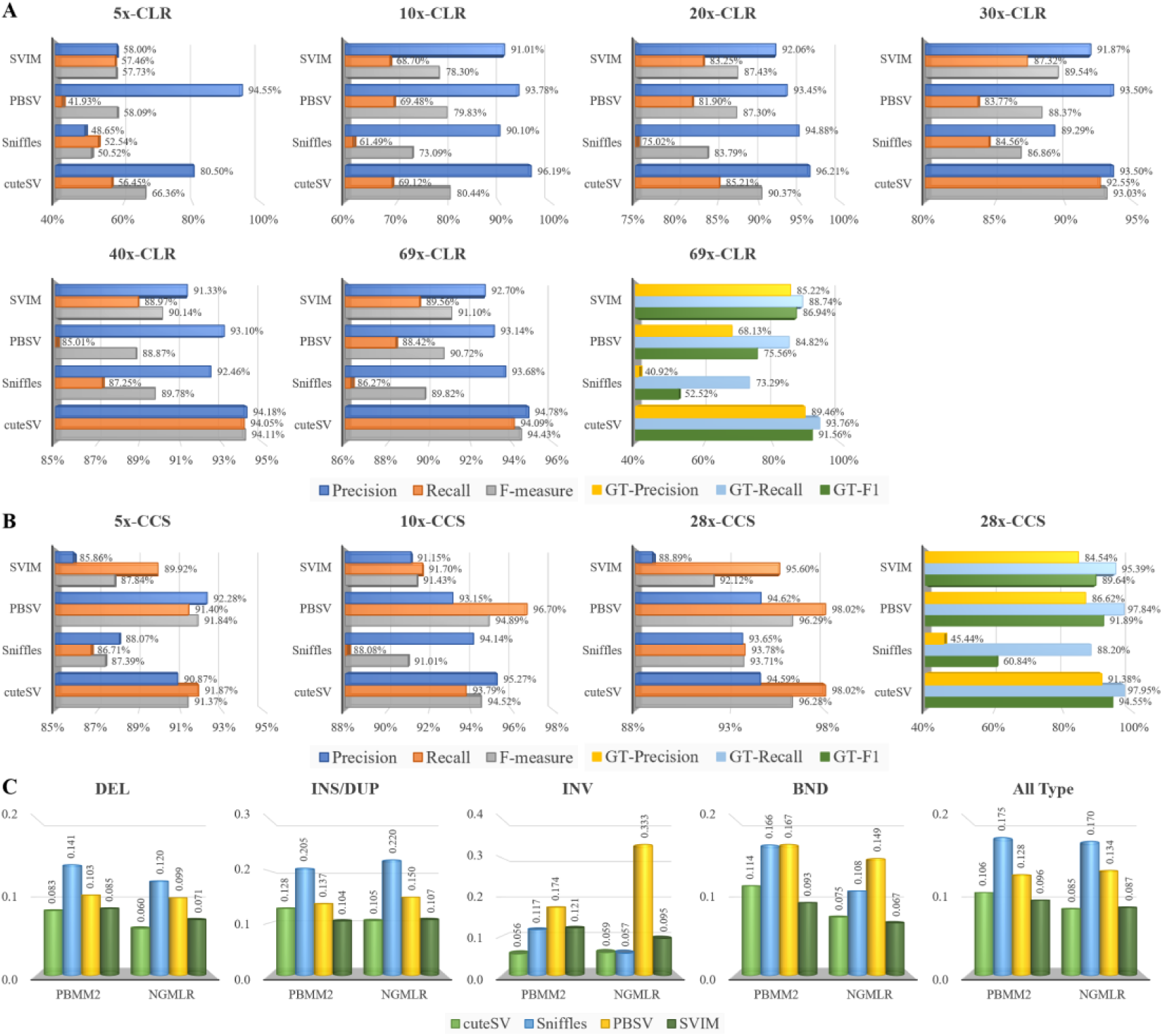
Benchmark results of the SV callers on HG002 PacBio CLR and CCS datasets. (**A**) Precisions, recalls and F-measures on the HG002 PacBio CLR datasets. (**B**) Precisions, recalls and F-measures on the HG002 PacBio CCS datasets. (**C**) Mendelian-Discordance-Rates (MDRs) on the GIAB Ashkenazi Trio PacBio CLR datasets for various SV types and long-read aligners. Worth noting that the lower the MDR is, the more accurate the corresponding SV callset is.

We further randomly down-sampled the original dataset to 5×, 10×, 20×, 30× and 40× to assess the ability of the SV callers on lower coverage datasets (Figure 2A and Supplementary Figure 1A). cuteSV almost kept achieving higher recalls and F1s on almost all datasets, indicating that it maintained outstanding sensitivity without loss of accuracy. It is worth noting that cuteSV simultaneously achieved >90% F1 and >86% GT-F1 at 20×. This is beneficial for practical use, since more cost-effective sequencing plans can be considered with this result. However, this is still hard for other callers since their recalls and F1s were much lower on the down-sampled datasets.

Besides PacBio CLR data, we also assessed the ability of the callers on the newly promoted PacBio CCS dataset of the same sample [39] [40] (coverage: 28×, mean read length: 13478 bp). The results of cuteSV and PBSV were very close to each other (Figure 2B) (precision: 94.6%, recall: 98.0%, F1: 96.3%), and they outperformed Sniffles and SVIM by 1% to 6% on various statistics. For genotyping assessment, cuteSV achieved the best performance on various GT-statistics, and outperformed the other three callers 2% to 34% on GT-F1.

We also randomly down-sampled the dataset to 5× and 10× to further assess the callers (Figure 2B and Supplementary Figure 1B). On the 10× down-sampled dataset, PBSV achieved the highest recall (96.70%) and cuteSV was the runner-up (93.79%). The slightly lower recall of cuteSV was mainly due to fact that the minimal signature size parameter setting of cuteSV (default value: 30bp) was longer than that of PBSV (default value: 20bp), which hindered cuteSV in identifying more SVs using evidence that was smaller in size. Moreover, it was also observed that 5× coverage was enough to achieve >90% precision, recall and F1 for both cuteSV and PBSV. These results suggest that with higher sequencing quality, data size requirements become smaller, so more economical sequencing strategies could be feasible in genomics studies.

We further used GIAB Ashkenazi Trio callsets (HG002, HG003 and HG004) to calculate the Mendelian-Discordance-Rate (MDR) of the long-read-based SV callers to more comprehensively assess their ability to detect various types of SVs. The results (Figure 2C) suggest that the MDRs of cuteSV and SVIM are lowest for almost all types of SV (i.e., All Type: 0.106 and 0.085 for cuteSV, and All Types: 0.096 and 0.087 for SVIM), indicating that the callsets produced by them are more plausible. The MDR of PBSV is higher but comparable (All Types: 0.128 and 0.134), while the MDR of Sniffles (All Types: 0.175 and 0.170) is much higher, indicating that its callset could be less accurate.

### SV detection with HG002 ONT PromethION data

We further assessed the yields of the SV callers on a newly published ONT PromethION dataset of the HG002 human sample (mean read length: 17335 bp, coverage: 47×, available at ftp://ftp.ncbi.nlm.nih.gov/giab/ftp/data/AshkenazimTrio/HG002_NA24385_son/UCSC_Ultralong_OxfordNanopore_Promethion/), which was aligned by using Minimap2. The benchmark results (Figure 3A) indicate that cuteSV achieved the highest precision (92.15%), recall (96.61%) and F1 (94.33%), and its outperformance of Sniffles and SVIM was more obvious than with the PacBio CLR dataset. It is also worth noting that PBSV crashed for this dataset. On the randomly down-sampled datasets (5×, 10× and 20×), the outperformance by cuteSV was remarkable as well. Particularly, cuteSV found most (85%) of the ground truth SVs only at 10× coverage with high precision (93.07%) and F1 (88.85%). However, at the same coverage, the F1s of Sniffles, SVIM and PBSV were at least 7% lower than that of cuteSV, indicating that they had lower yields. As for the genotyping (Figure 3B and Supplementary Figure 1C), cuteSV still achieved better performance and surpassed others 9% at least on GT-F1. This indicates that the genotyping of cuteSV could be more accurate.

**Figure 3.**
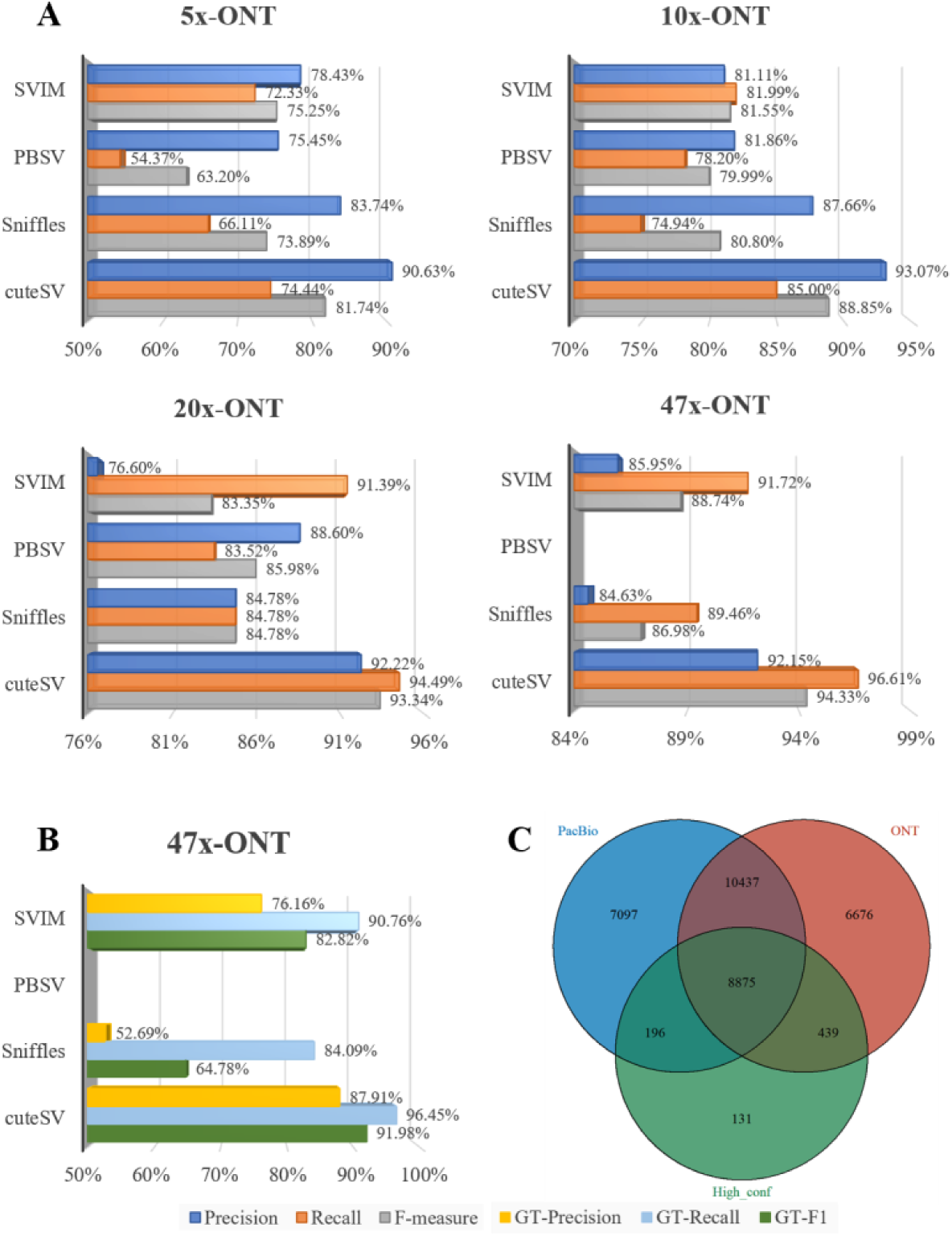
Benchmark results of the SV callers on the HG002 ONT PromethION dataset. **(A)** Precisions, recalls and F-measures on the HG002 ONT dataset and its down-sampling data. (**B**) GT-Precisions, GT-recalls and GT-F1 on the HG002 ONT dataset. (**C**) The Venn diagram of SV calls produced by cuteSV from HG002 PacBio CLR and ONT PromethION datasets (indicated by “PacBio” and “ONT”, respectively), and the SV ground truth callset of the HG002 sample made by GIAB (indicated by “High_conf”).

We further compared the SV callsets produced by cuteSV from the HG002 PacBio CLR and ONT datasets, respectively. There are 26605 and 26427 SVs in the PacBio and ONT callsets, respectively (see Supplementary Table 1). 7293 SVs (27.41%) of the PacBio callset and 7115 SVs (26.92%) of the ONT callset are unique. Moreover, of the 7293 PacBio-only calls, 54.15% (3949 of 7293) and 21.20% (1546 of 7293) are respectively insertions and deletions, and of the 7115 ONT-only calls, the corresponding numbers are respectively 47.94% (3411 of 7115) and 41.00% (2917 of 7115). Previous studies indicate that there are usually more false positive insertion calls made from PacBio datasets [23], and more false positive deletions made from ONT datasets [24]. So, the fractions of the unique calls of the PacBio and ONT callsets are plausible. Moreover, it is also observed that there are some SVs only being discovered in the ONT dataset mainly because of its larger read lengths. An example is shown in Supplementary Figure 2. A 6481 bp insertion (breakpoint at chr1:9683994) was only detected in the ONT reads, as the ONT reads spanning this event are longer and a large proportion of them carry significant insertion signals in their CIGARs. However, none of the PacBio reads aligned to this locus have strong SV signals, possibly because they are shorter and the aligners cannot produce alignments with such large insertions. Moreover, there were 33751 distinctive SV calls in the PacBio- and ONT-based callsets and they covered 98.64% (9510 of 9641) of the ground truth callset of HG002 (Figure 3C) (i.e., they had a higher sensitivity than all the callsets made by the SV callers with a single dataset). This suggests that combining multiple datasets to produce a high-quality SV callset is feasible.

### The performance of the SV callers

We assessed the speed (Figure 4A) and memory footprint (Figure 4B) of the SV callers on the 69× HG002 CLR dataset. The speeds of the benchmarked approaches with a single CPU thread were comparable: Sniffles was the fastest, taking 367 minutes, followed by SVIM (991 minutes), cuteSV (1279 minutes) and PBSV (1715 minutes). The relatively slower speed of cuteSV was mainly because it uses multiple methods to extract signatures of SVs and perform its high-quality genotyping. Further, we asked the SV callers to run with multiple CPU threads, but it was observed that neither Sniffles nor PBSV had an obvious speedup with more CPU threads, and SVIM does not support multiple-threading computing. This is possibly due to their implementations. However, cuteSV showed a nearly linear speedup with the number of CPU threads, and the wall clock time was greatly reduced. For example, with 16 CPU threads, cuteSV used only 104 minutes to process the whole dataset. Moreover, the memory footprint of cuteSV (0.38GB) was smaller than that of other approaches by two orders of magnitude (i.e., Sniffles: 17.61GB, PBSV: 10.57GB, SVIM: 19.10GB). With its quasi linear multiple-threads speedup and low memory footprint, we realized that cuteSV is a highly scalable SV detection tool, which is suited to high performance computing platforms and large-scale data analysis tasks such as SV detection in many samples.

**Figure 4.**
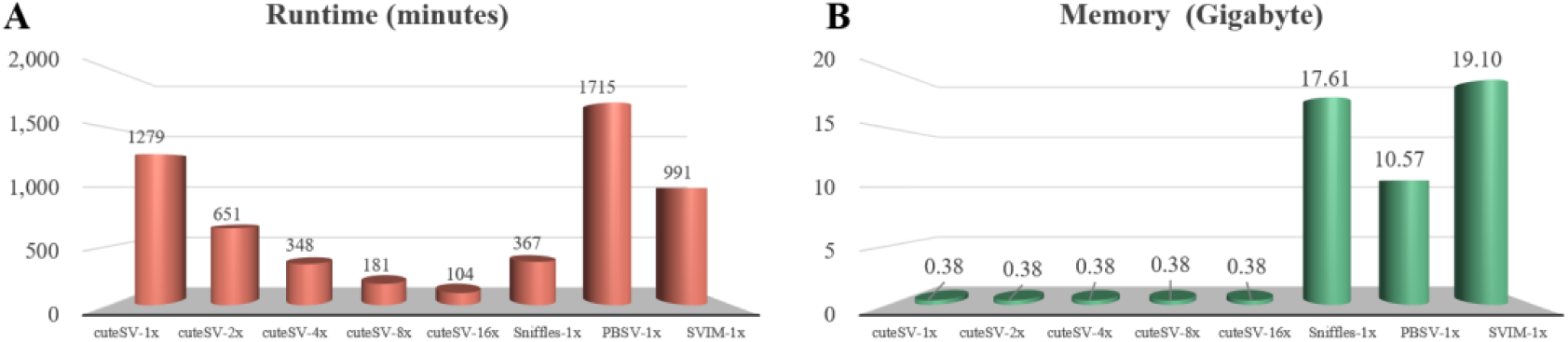
Performance of the benchmarked SV callers. (**A**) The runtime and (B) the memory footprints of the SV callers are shown. The results of cuteSV with various numbers of CPU threads are given, and for other SV callers, only the results using a single CPU thread are shown as the results of Sniffles and PBSV with multiple CPU threads are quite similar to that of a single CPU and SVIM does not support multiple thread computing.

### The effects of various read aligners on SV calling

To investigate the effects of various aligners on SV detection, we employed two aligners, PBMM2 and NGMLR, to separately align the reads of the 69× HG002 PacBio CLR dataset, and used their alignments as inputs to the four SV callers to produce various callsets. The callsets were then evaluated, and the effects of the aligners on the sensitivity and accuracy of SV detection were observed. Some details are as follows.

1. We collected SV calls marked as false-negative (FN) by Truvari and compared them to investigate the effect of the two aligners on the sensitivity of the SV callers. The differences between the callsets produced by same SV callers with the alignments of different aligners are shown in Figure 5A and Supplementary Table 2. The results indicate that, overall, the SV callers had higher sensitivities (fewer FN calls) on insertions and deletions of less than 1,000 bp with the alignments produced by PBMM2. Moreover, PBSV discovered 436 more insertions with PBMM2 than with NGMLR, the lengths of which are between 200 bp and 700 bp, and 369 of which coincided with the sequences of the Alu family. This issue is worth noting since MEI is a major category of SVs. On the other hand, cuteSV, Sniffles, PBSV and SVIM respectively discovered 109, 283,127 and 113 more larger SVs (size >1000 bp) with the alignments of NGMLR compared to their own callsets based on the alignments of PBMM2, which is also worth noting.
2. We further investigated the false-positive (FP) calls of the SV callers based on various aligners (see Figure 5B and Supplementary Table 3). The callsets of cuteSV, Sniffles and PBSV with PBMM2 had slightly higher numbers (155, 54 and 43, respectively) of <1000 bp FP insertions and deletions than those with NGMLR. While the callset of SVIM with NGMLR had a higher number (45) of FP calls. For >1000 bp SVs, the numbers of FPs were very close for the corresponding callsets of cuteSV, PBSV and SVIM, but there were 32 more FPs in the callset of Sniffles with NGMLR than that with PBMM2.
3. The MDRs of the callsets with various aligners were also assessed (Figure 5C). It was observed that all four callers had decreased MDRs (by 0.35% to 5.83%) on deletions and translocations, indicating that NGMLR kept better Mendelian consistency for these types of SVs. While no significant difference was observed for insertions/duplications and inversions with the two aligners. Moreover, only PBSV had obviously higher overall MDRs with NGMLR, indicating that PBMM2 could be a better choice for it.

**Figure 5.**
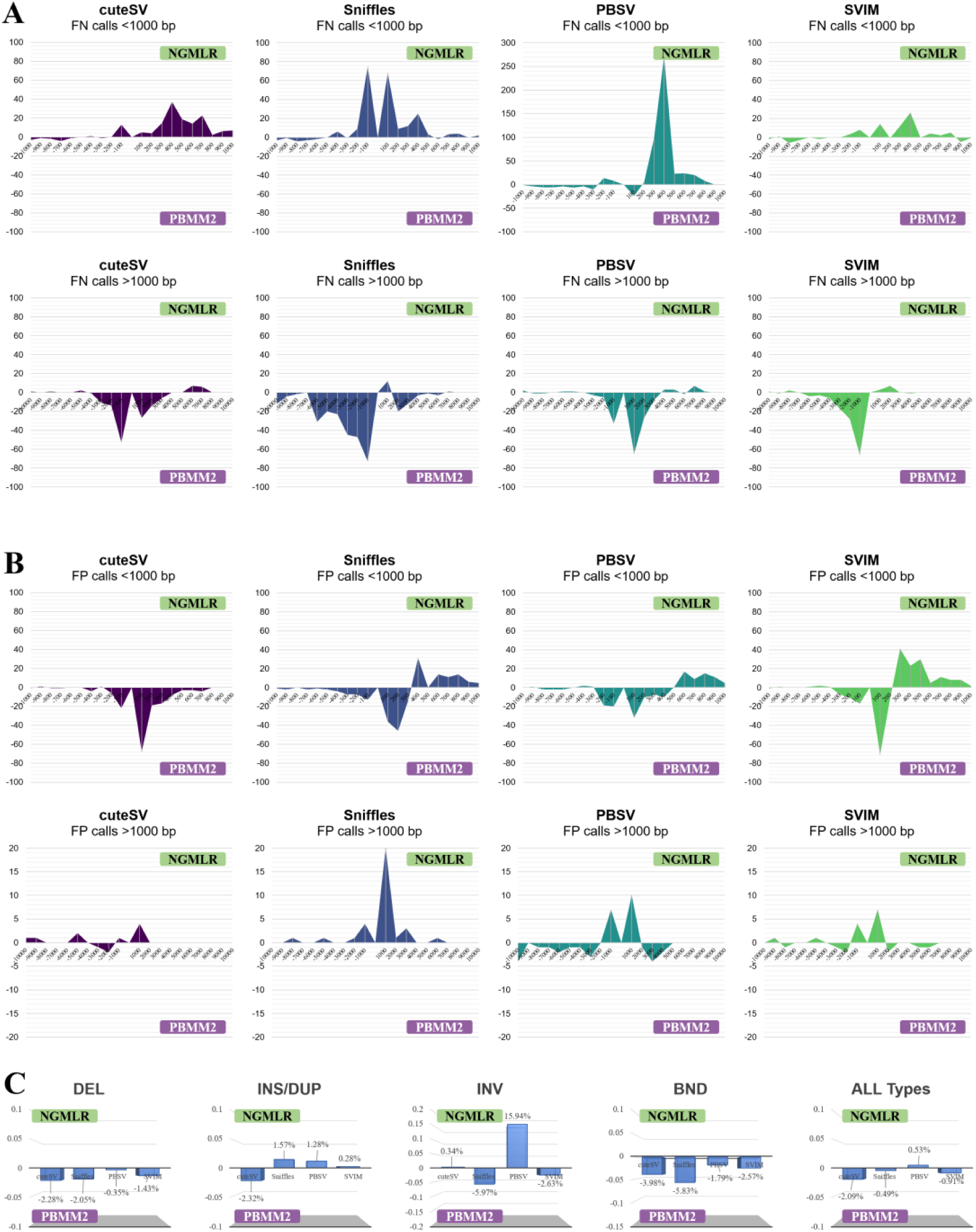
Comparison of the callsets of the PacBio CLR dataset produced with various long-read aligners. (**A**) The difference between the false-negative (FN) calls in various SV sizes with NGMLR and PBMM2. (**B**) The difference between the false-positive (FP) calls in various SV sizes with NGMLR and PBMM2. (**C**) The different MDRs between NGMLR and PBMM2. It is worth noting that a positive value indicates the occurrence of more corresponding calls (e.g., FN, FP and MDR) when using NGMLR, whereas a negative value means the generation of more corresponding calls with PBMM2. The smaller the overall difference, the better the compatibility with the long-read aligners.

### The results of cuteSV with various configurations of parameters

We assessed the yields of cuteSV with various configurations of two critical parameters, –min_support and –min_size (-s and -l in the software, respectively), on various coverages (5×, 10×, 20×, 30×, 40× and 69×) of HG002 CLR datasets.

The –min_support parameter indicates the minimal number of supporting reads to make an SV call. As the results displayed in Supplementary Figure 3A and Supplementary Table 4 show, with the default setting of –min_size (–min_size = 30), cuteSV achieved the best yields when –min_support was configured at 1 to 10 for the various coverages, and there is an obvious trade-off between precision and recall (i.e., setting –min_support to smaller numbers might result in higher sensitivity but lower precision, and vice versa).

The –min_size parameter indicates the minimal size of SV signature considered in clustering. Keeping the –min_support parameter fixed, we investigated the results of cuteSV with two different settings for –min_size (i.e., –min_size = 30 and –min_size = 50, Supplementary Figure 3B). At various coverages, the accuracies of cuteSV were 0.12% to 1.38% higher with the setting of –min_size = 50, while the recall rates were -0.06% to 1.09% higher with –min_size = 30. This indicates that setting –min_size with smaller numbers might result in higher sensitivity but lower accuracy, and vice versa. It is worth noting that, although the trade-off exists, for each of the coverages, the F1s of cuteSV with various settings are quite close to each other (the difference is less than 1%).

### SV detection with NA19240 PacBio CLR data

A PacBio CLR dataset from another well-studied human sample (NA19240) [41] was also employed (mean read length: 6503 bp, coverage: 40×) to benchmark the SV callers more comprehensively. A callset for this sample [42] was used as the ground truth.

The precision, recall and F1-measure of the benchmarked SV callers are shown in Supplementary Figure 4 and Supplementary Table 5. cuteSV almost had the highest F1 rates for all types of SVs (i.e., DEL: 62.79%, INS/DUP: 54.28%, INV: 17.36% and All Types: 57.66%). SVIM achieved better recall rates for all types of SVs and outperformed cuteSV 1.9% on All Types, which is mainly due to its larger number of predictions (total call: 31053 vs. 27294), however, its F1s were similarity to that of cuteSV mainly due to its relatively poor accuracies. Similarly, Sniffles and PBSV respectively obtained 0.35% and 3.22% higher precision than cuteSV, whereas their lower recall rates decreased the F1s of them. Overall, cuteSV achieved the comparable performance for this dataset.

## Discussion

Long-read sequencing technologies provide opportunities for comprehensive SV discovery. However, utilizing the advantage of long reads is still non-trivial due to complicated alignments. Herein, we propose cuteSV, a sensitive, fast and scalable long-read-based SV detection approach. We show how to use a series of signature extraction methods and a tailored clustering-and-refinement method to implement a precise analysis of SV signatures to achieve good yields and performance simultaneously. The cuteSV approach has four major contributions as follows.

1. Read aligners could produce fragile alignments around SV events, for example, some reads could have several smaller insertions and/or deletions in their alignments around a larger SV event. Such alignments usually mislead SV detection approaches to make incorrect calls. However, using specifically designed methods, cuteSV can adaptively combine the chimeric alignments in reads to effectively recover the signatures of the real SV events. An example in Supplementary Figure 5 shows how a 272bp insertion event was successfully discovered by cuteSV, while none of the other benchmarked SV callers managed to do so due to fragile alignments.
2. There are multi-allelic heterozygous SVs in some local regions, so it is even more difficult when the multiple alternative alleles have the same SV type. Since most state-of-the-art SV callers rely on genomic-position-based clustering of chimeric reads, reads spanning various SVs in a local region could be binned in one cluster, and then incorrect SV calls could be made. cuteSV solves this problem by using a tailored cluster refinement approach, which precisely analyzes the SV signatures of the reads in the same cluster. This operation helps to distinguish reads spanning different SV alleles, and correctly bins them by various sub-clusters, which greatly helps the detection of such complex SV events. Two examples are shown in Supplementary Figure 6. One is a heterozygous insertion event (a 180bp and a 36bp insertion in the same region) and the other is a heterozygous deletion event (a 123bp and a 37 bp deletion in the same region). Only cuteSV and PBSV successfully discovered them in the benchmark.
3. With its tailored signature extraction and clustering methods, cuteSV enables more sensitive detection and more accurate genotyping of SVs. The advantage is more obvious for relatively low coverage datasets. The benchmark results suggest that cuteSV can discover most SVs in 20× coverage datasets for human samples without loss of accuracy, and the performance of genotyping could be consistent with its yields. This is helps to make more flexible sequencing plans in large-scale genomics studies (e.g., population genomics studies) and to achieve goals in a more economical way.
4. Most state-of-the-art SV callers are still not very scalable (i.e., they cannot fully take advantage of computational resources for speedup). This could be problematic when using large datasets since more computational nodes would be needed and/or the time cost might be very high. However, cuteSV has good scalability with its block division implementation, which enables it to process the input dataset in a parallel way and achieve a nearly linear speedup with the number of CPU threads. This makes it very suited to modern HPC resources and helpful for upcoming large-scale genomics studies.

The benchmark results also indicate that there are still some SVs that cannot be successfully detected by cuteSV. We investigated the intermediate results of cuteSV and found that most of the false negative calls were due to the read alignments being inaccurate or not informative enough. Two typical examples are shown in Supplementary Figures 7 and 8. In Supplementary Figure 7, a 98bp deletion case is shown in which deletion signatures emerge in nearly all the reads around the event. However, the size of the deletions in the alignment are not correct (i.e., most of them are around 50bp). In Supplementary Figure 8, a 707bp insertion case is shown in which insertion signatures also emerge and their sizes are close to the SV event, but the positions of the breakpoints in the reads are quite far from the ground truth breakpoint. Under such circumstances, all the benchmarked SV callers made SV calls, but the sizes and/or positions are incorrect since the read alignments are misleading. We therefore realize that important future work could be done to develop more accurate long read aligners.

## Conclusion

Long-read sequencing technologies provide the opportunity to build a comprehensive map of structural variations of the human genome. However, there are still technical issues to be addressed to further improve the sensitivity, accuracy and performance of long-read-based SV detection, and the development of advanced computational approaches is in demand.

In this article, we provide a novel long-read-based SV detection approach, cuteSV. It enables the thorough analysis of the complex signatures of SVs implied by read alignments. Benchmark results demonstrate that cuteSV achieves good yields and performance simultaneously. Moreover, it has an outstanding ability to detect SVs with low coverage sequencing data, and it also has high scalability for handling large-scale datasets. We believe that cuteSV has great potential for cutting-edge genomics studies.

## Materials and methods

### Read alignment

cuteSV uses sorted BAM files as input, and is supported by employing state-of-the-art long-read aligners to compose SV detection pipelines. Aligners with a good ability to handle large insertions/deletions in reads and/or produce accurate split alignments are preferred, since cuteSV extracts important SV signatures from such alignments. In the Results section, it is demonstrated that state-of-the-art aligners such as PBMM2 and NGMLR are suitable for the cuteSV approach.

### Extraction of SV signatures implied by CIGARs

Given a set of aligned reads, cuteSV separately analyzes the detailed alignment of each read. Mainly, it extracts two categories of SV signatures, namely the long insertions/deletions in CIGARs and split alignments, respectively. In the cuteSV approach, the signatures of SVs are represented as a 3-tuple (i.e., *SIG* = (*pos, len, ID*), where *pos* indicates the starting position on the reference genome, *len* indicates the size of SV and *ID* indicates the unique read ID). cuteSV composes the SV signals implied by read alignments into *SIGs*, and *SIGs* of various reads are clustered and analyzed to call SVs.

cuteSV uses a precise method to extract and merge the long insertions/deletions in CIGARs and to transform them into SV signatures. This method enables the recovery of SV signatures of long insertions/deletions which are initially partitioned as multiple trivial indels by read aligners, and it facilitates the inference of the breakpoints and lengths of SVs. More specifically, cuteSV extracts insertions/deletions >30 bp in size as described by the CIGARs of the reads, and composes them into *SIGs* with their positions, lengths and read IDs. For two signatures, *SIG*_1_ and *SIG*_2_, cuteSV merges them if they meet the following condition:

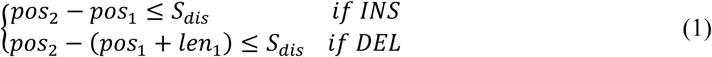

where *S*_*dis*_ is a threshold of the distance between the two signatures (default value: 500 bp). The two signatures are then merged as *SIG*_*M*_ = (*pos*_1_, *len*_1_ + *len*_2_, *ID*). The merging operation enhances the signatures of large insertions/deletions, and all the remaining signatures after merging are used as informative signatures for further processing.

### Extraction of SV signatures implied by split alignments

For a read with split alignment(s) (described by its primary and supplementary alignments), cuteSV records each split by a 6-tuple (also termed as a “segment”), *Seg* = (*read*_*s*_, *read*_*e*_, *Ref*_*s*_, *Ref*_*e*_, *Chr, Strand*), where *read*_*s*_, *read*_*e*_, *Ref*_*s*_, *Ref*_*e*_ respectively indicate the starting and end coordinates on the read and reference genome, and *Chr* and *Strand* respectively indicate its chromosome and orientation. cuteSV uses several heuristic rules (detailed below) to recover SV signatures from these *Segs*.

1. Extraction of deletion/insertion signatures. If two segments, *Seg*_1_ and *Seg*_2_, are adjacent on the read and aligned to the same chromosome with identical orientations, cuteSV computes *Diff*_*dis*_ = (*Ref*_2*s*_ − *Ref*_1*e*_) − (*read*_2*s*_ − *read*_1*e*_) and *Diff*_*olp*_ = *Ref*_1*e*_ − *Ref*_2*s*_. If *Diff*_*olp*_ < 30 *bp* and *Diff*_*dis*_ ≥ 30 *bp*, cuteSV considers that the two segments indicate a deletion event, and composes a deletion signature:

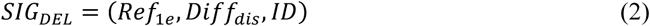 Moreover, if *Diff*_*olp*_ < 30 *bp* and *Diff*_*dis*_ ≤ −30 *bp*, cuteSV considers this an insertion event and composes an insertion signature:

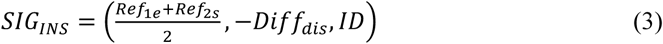
2. Extraction of duplication signatures. If two adjacent segments are mapped to similar positions (i.e., *Diff*_*olp*_ ≥ 30 *bp*), cuteSV composes a duplication signature:

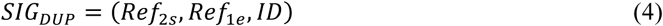
3. Extraction of inversion signatures. If two adjacent segments are mapped to the same chromosome but to different strands, cuteSV composes an inversion signature:

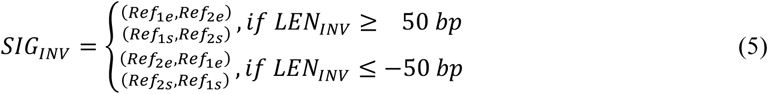

where *LEN*_*INV*_ = *Ref*_2*e*_ − *Ref*_1*e*_ indicates the length of the event (with direction). It is also worth noting that the primary strand of the inversion is unknown, so cuteSV records all possible situations.
4. Extraction of translocation signatures. If two adjacent segments are mapped to different chromosomes, and the two segments are <100bp distant on the reads, cuteSV composes a translocation signature:

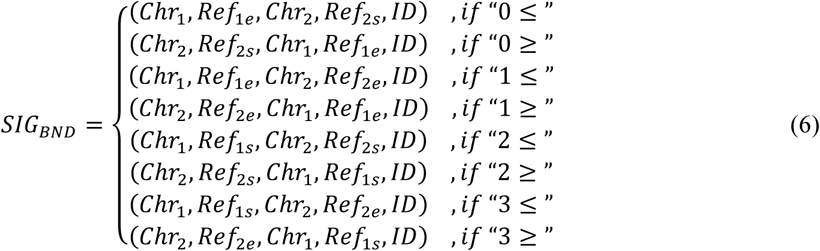

where 0, 1, 2, 3 indicate the combination of strands “++”, “+-”, “-+”, “--”, respectively, and “≤” and “≥” respectively mean the chromosome ID of *Chr*_1_ is in front of that of *Chr*_2_ and vice versa.
5. cuteSV uses a specifically designed method to extract the signatures of a complex kind of SV, when there is a mobile insertion in between two duplicated sequences. An example is shown in Supplementary Figure 9, in which there are two duplicated local sequences (both of their mapped positions are around Chr1:73594981), and there is another local sequence within them in the read (whose mapped position is in a decoy sequence of hs37d5). cuteSV extracts a series of signatures, including a duplication (as shown in Eq.4), a translocation (as shown in Eq.6) and an insertion (as shown in Eq.7), to describe such complex SV events.

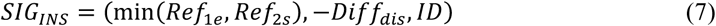

### Clustering of SV signatures

cuteSV clusters the collected SV signatures (*SIG*s) in a two-step approach. In the first step it clusters the signatures by their genomic positions and types in order to bin the signatures in various local regions, and in the second step it clusters the signatures by their length in order to distinguish the signatures in similar regions but belonging to heterozygous SVs (i.e., in such loci, there are different but similar SVs in both of the two alleles). Some details are as follows.

In the first step, cuteSV sorts all the SV signatures by their genomic positions and types (i.e., deletions, insertions, duplications, inversions and translocations). For each category, cuteSV initially creates a new cluster, and scans all the signatures from upstream to downstream and adds them into the cluster using an iterative approach. More precisely, for a newly-scanned SV signature *SIG*_*i*_, cuteSV adds it into the cluster if there is at least one signature *SIG*_*j*_ in the cluster which meets the following condition:

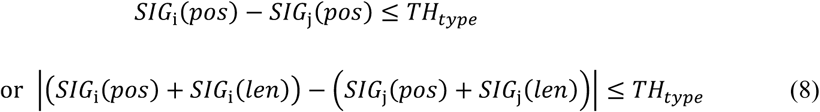

where *TH*_*type*_ is a threshold of the distances among the clustered signatures, and different values are used for various types of SVs (*TH*_*type*_ is typically configured to between 50 and 500 bp). If *SIG*_*i*_ cannot be added into the cluster, cuteSV creates a new cluster only having *SIG*_*i*_ and goes to the next SV signature.

In the second step, cuteSV initially checks the numbers of signatures of the generated clusters and discards the clusters with too few signatures. The remaining clusters are then refined by the following methods according to their SV type.

1. Refinement of deletion/insertion clusters. Given a deletion or insertion cluster, cuteSV sorts all its signatures by their sizes, and computes a parameter, *Len*_*bias*_, with the following equation:

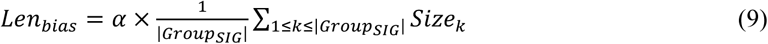

where *α* is a weighting parameter, |*Group*_*SIG*_| is the number of the signatures in the cluster, and *Size*_*k*_ is the size of the k-th longest signature in *Group*_*SIG*_. The configuration of *α* is related to the SV type. The default value of *α* is 0.2 for an insertion cluster and high error rate reads (e.g., PacBio CLR and ONT reads), *α* = 0.65 for low error rate reads (e.g., PacBio CCS reads) and *α* = 0.3 for a deletion cluster (regardless of error rate). With *Len*_*bias*_, cuteSV divides the cluster into sub-clusters of which each is a potential SV allele in a local genomic region. More specifically, cuteSV initially creates one sub-cluster and add the signatures with the largest size into the sub-cluster. It iteratively scans the signatures by size (from largest to smallest). For instance, if the difference between the size of a newly-scanned signature and that of the last signature added to the sub-cluster is smaller than *Len*_*bias*_, cuteSV adds the newly-scanned one into the sub-cluster. Otherwise, a new sub-cluster is created with the newly-scanned signature. cuteSV recognizes the generated sub-clusters with the highest number of signatures as a “major allele” sub-cluster if it meets the following conditions:

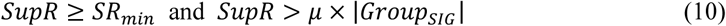

where *SupR* and *SR*_*min*_ are the number of its supporting reads and the threshold of the minimum number of supporting reads, respectively. *μ* is a weighting parameter, and its default values for an insertion cluster are 0.6 and 0.65 when using PacBio CLR/ONT reads and PacBio CCS reads, respectively. For a deletion cluster, the default values are 0.7 and 0.35 for the datasets mentioned above, respectively. If a “major allele” sub-cluster exists, cuteSV recognizes each of the remaining sub-clusters as a “minor allele” sub-cluster if it has more than *SR*_*min*_ signatures. However, there is occasionally no “major allele” sub-cluster existing due to lack of enough supporting reads. cuteSV uses another heuristic rule in which it recognizes the two largest sub-clusters *SupR*_*first*_ and *SupR*_*second*_ as “major allele” and “minor allele” sub-clusters if they meet the following conditions:

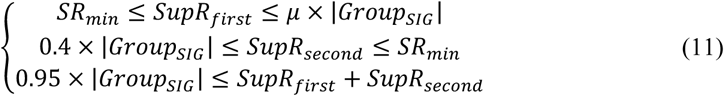 This rule indicates that almost all of the SV signatures support the two alleles. Meanwhile, both of them occupy >40% of the supporting signatures of the cluster.
2. Refinement of duplication/inversion clusters. Given a duplication or inversion cluster, cuteSV initially creates one or more sub-clusters such that the breakpoints of all the signatures in the same sub-cluster are within 500bp. cuteSV recognizes the “major allele” and “minor allele” sub-clusters with a heuristic rule similar to Eq.10 but respectively sets *μ* = 1/3 for duplication and inversion.
3. Refinement of translocation clusters. Given a translocation cluster, cuteSV initially creates one or more sub-clusters such that the breakpoints of all the signatures in the same sub-cluster are within 50bp. Furthermore, cuteSV recognizes the “major allele” and “minor allele” sub-clusters with a heuristic rule similar to Eq.10 but sets *μ* = 0.6 and *SR*_*min*_ as half of the value used for deletion/insertion clusters. This setting mainly considers the diverse combinations of chromosomes and orientations of translocation events and is beneficial for sensitivity.

### SV calling and genotyping

For each cluster of signatures, cuteSV computes the weighted average of the positions and sizes to predict the breakpoint(s) and size of the corresponding SV, and removes the predicted SVs of <30 bp in size.

Moreover, cuteSV supports the implementation of SV genotyping (which is an optional step in the tool). cuteSV uses a local genomic region for a predicted SV to analyze the ratio of supporting reads, which is defined as:

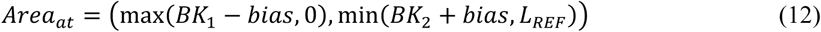

where *BK*_1_ is the first breakpoint, *BK*_2_ is the next breakpoint, *bias* means the scope of extension which is equal to *TH*_*type*_ and *L*_*REF*_ indicates the maximum length of chromosome taken into consideration. cuteSV counts the numbers of all the reads supporting the reference allele and the reads supporting the predicted SV within *Area*_*at*_. Various genotypes are assigned by cuteSV according to the ratio of supporting reads (i.e., homozygous reference allele (0/0) for a ratio ≤ 0.2, homozygous variant allele (1/1) for a ratio ≥ 0.8, and heterozygous variant allele (0/1) for a ratio in between).

### Implementation of long-read mapping and SV calling

We used PBMM2 (version 1.0.0), NGMLR (version 0.2.3) and Minimap2 (version 2.17) to implement the read alignment of the benchmarking datasets (Supplementary Table 6). The parameter “--preset” of long-read aligners was tuned for different sequencing technologies. Samtools (version 1.9) was employed for read extraction, sorted BAM generation and sequencing data down-sampling.

Sniffles (version 1.0.11), PBSV (version 2.2.0), SVIM (version 0.4.3) and cuteSV (version 1.0.1) were benchmarked with the sorted BAM files as input. Specifically, for Sniffles, the configuration “-l 50 -s 2/3/4/4/5/10” was used for PacBio CLR datasets, “-l 30 -s 1/2/3” for was used for PacBio CCS reads, and “-l 30 -s 2/3/4/10” was used for ONT PromethION reads. For PBSV, default settings were used for PacBio CLR and ONT PromethION data, and “--preset CCS” was used for PacBio CCS data. For SVIM, the configuration “--min_sv_size 30” was employed for all datasets in this study. For cuteSV, the configuration “-l 30 --max_cluster_bias_INS 100 --diff_ratio_merging_INS 0.2 --diff_ratio_filtering_INS 0.6 --diff_ratio_filtering_DEL0.7” was used for PacBio CLR (“-s 2/3/4/4/5/10”) and ONT PromethION reads (“-s 2/3/4/10”), and “-l 30 --max_cluster_bias_INS 200 --diff_ratio_merging_INS 0.65 --diff_ratio_filtering_INS 0.65 --diff_ratio_filtering_DEL0.35 -s 1/2/3” was used for PacBio CCS reads.

The used command lines for the tools are in the Supplementary Notes.

### Evaluation of SV callsets

The evaluation of the HG002 human sample is run with Truvari (version 1.1), and the high confidence insertion and deletion callsets (version 0.6) are used as gold standard sets. Before evaluation, we preprocess the SV calls of each tool. For Sniffles, we discard inversions and translocations, and transform duplications to insertions. For SVIM, we delete SV calls with a quality score of less than 40, as the author recommends, and transform duplications to insertions as well. For PBSV and cuteSV, we only select insertions and deletions for assessment. Then, BGZIP and TABIX are employed to compress and index the processed VCF files.

Only SV calls between 50 bp and 10 kbp in size that are within the GIAB high confidence regions (version 0.6) are considered for evaluation. A prediction is determined as a true-positive (TP) when meeting the following conditions:

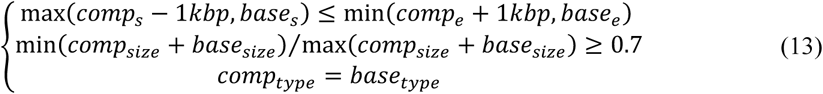

where *comp*_*s*_, *comp*_*e*_, *comp*_*size*_ and *comp*_*type*_ indicate start coordinate, stop coordinate, size and SV class of a prediction, and *base*_*s*_, *base*_*e*_, *base*_*size*_ and *base*_*type*_ are starting coordinate, stopping coordinate, size and SV class of a call in the truth set, respectively. On the other hand, a prediction is determined as a false-positive (FP) if it does not satisfy Eq.13. A false-negative (FN) is assigned when there is a base call that cannot be detected by any SV calls.

Based on the results above, Precision (or the ratio of TPs to total calls in predictions) is defined as

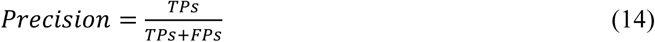

Similarly, Recall (or the ratio of TPs to total calls in the truth set) is defined as

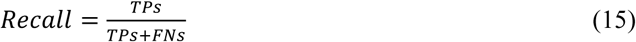

F-measure (F1 score), a measurement of weighted averaging of both precision and recall, is defined as

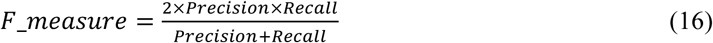

The evaluation of the NA19240 is similar to the HG002 benchmarking. We adopt Eq.13 to assess every deletion, insertion (duplication regarded as a subset of insertion) and inversion against the callsets generated from the study [42], and Eqs.14 to 16 are applied for summarizing the performance of SV calling.

For the evaluation of the Ashkenazi trio, the callsets of parents (i.e., HG003 and HG004) produced by corresponding methods are adopted as truth sets in order to measure the performance of an SV caller via the pedigree between offspring and parents. For each SV call of a son (i.e., HG002) except translocations, we use Eq.13 to assign true-positive calls. As an arbitrary rearrangement type, a translocation or BND is considered precise when meeting the following conditions:

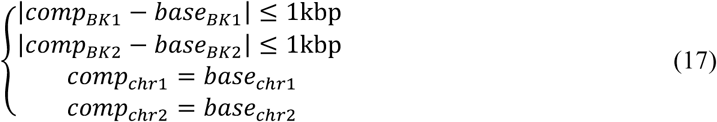

where *BK*1, *BK*2, *chr*1 and *chr*2 are the combination of breakends and chromosomes of a call on the offspring and its parents, respectively. Therefore, the ratio of MDR can be reckoned via

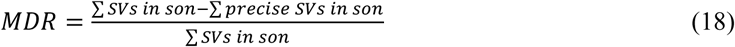

To evaluate the computational performance of the four SV callers under different threads, runtime and memory usage are assessed using the “/usr/bin/time -v” command of the Linux Operating System. In the output results of the command, “Elapsed (wall clock) time” and “Maximum resident set size” indicate the elapsed runtime and memory consumption, respectively. It is worth noting that because SV calling performed by PBSV involves two steps (i.e., discover and call), we use the sum of the wall clock time of both steps as the final elapsed runtime, whereas the memory consumption depends on the maximum memory usage of the two runs.

For more details please see the Supplementary Notes.

## Authors’ contributions

T.J. conceived and supervised the project. T.J. developed cuteSV. T.J., B.L. and Y.J. performed the analysis. T.J., B.L and J.L. wrote the manuscript. All authors read and approved the final manuscript.

## Availability of data and material

The human reference genome, raw sequencing data, alignments and gold standard truth callsets used in this study are available from the respective publications and websites listed in Supplementary Table 6. The implementation of cuteSV can be downloaded from GitHub (https://github.com/tjiangHIT/cuteSV).

## Ethics approval and consent to participate

Not applicable.

## Consent for publication

Not applicable.

## References

1. Sudmant PH, Rausch T, Gardner EJ, Handsaker RE, Abyzov A, Huddleston J, Zhang Y, Ye K, Jun G, Fritz MH, et al: An integrated map of structural variation in 2,504 human genomes. Nature 2015, 526: 75–81.

2. Alkan C, Coe BP, Eichler EE: Genome structural variation discovery and genotyping. Nat Rev Genet 2011, 12: 363–376.

3. Rovelet-Lecrux A, Hannequin D, Raux G, Le Meur N, Laquerriere A, Vital A, Dumanchin C, Feuillette S, Brice A, Vercelletto M, et al: APP locus duplication causes autosomal dominant early-onset Alzheimer disease with cerebral amyloid angiopathy. Nature Genetics 2006, 38: 24–26.

4. Hedges DJ, Hamilton-Nelson KL, Sacharow SJ, Nations L, Beecham GW, Kozhekbaeva ZM, Butler BL, Cukier HN, Whitehead PL, Ma DQ, et al: Evidence of novel fine-scale structural variation at autism spectrum disorder candidate loci. Molecular Autism 2012, 3.

5. Weischenfeldt J, Symmons O, Spitz F, Korbel JO: Phenotypic impact of genomic structural variation: insights from and for human disease. Nat Rev Genet 2013, 14: 125–138.

6. Macintyre G, Ylstra B, Brenton JD: Sequencing Structural Variants in Cancer for Precision Therapeutics. Trends in Genetics 2016, 32: 530–542.

7. Dennenmoser S, Sedlazeck FJ, Iwaszkiewicz E, Li XY, Altmuller J, Nolte AW: Copy number increases of transposable elements and protein-coding genes in an invasive fish of hybrid origin. Mol Ecol 2017, 26: 4712–4724.

8. Lupski JR: Structural variation mutagenesis of the human genome: Impact on disease and evolution. Environ Mol Mutagen 2015, 56: 419–436.

9. Chiang C, Scott AJ, Davis JR, Tsang EK, Li X, Kim Y, Hadzic T, Damani FN, Ganel L, Consortium GT, et al: The impact of structural variation on human gene expression. Nat Genet 2017, 49: 692–699.

10. Zichner T, Garfield DA, Rausch T, Stutz AM, Cannavo E, Braun M, Furlong EEM, Korbel JO: Impact of genomic structural variation in Drosophila melanogaster based on population-scale sequencing. Genome Research 2013, 23: 568–579.

11. Jeffares DC, Jolly C, Hoti M, Speed D, Shaw L, Rallis C, Balloux F, Dessimoz C, Bahler J, Sedlazeck FJ: Transient structural variations have strong effects on quantitative traits and reproductive isolation in fission yeast. Nature Communications 2017, 8.

12. Levy S, Sutton G, Ng PC, Feuk L, Halpern AL, Walenz BP, Axelrod N, Huang J, Kirkness EF, Denisov G, et al: The diploid genome sequence of an individual human. PLoS Biol 2007, 5: e254.

13. Wheeler DA, Srinivasan M, Egholm M, Shen Y, Chen L, McGuire A, He W, Chen YJ, Makhijani V, Roth GT, et al: The complete genome of an individual by massively parallel DNA sequencing. Nature 2008, 452: 872–876.

14. Yoon ST, Xuan ZY, Makarov V, Ye K, Sebat J: Sensitive and accurate detection of copy number variants using read depth of coverage. Genome Research 2009, 19: 1586–1592.

15. Chen K, Wallis JW, McLellan MD, Larson DE, Kalicki JM, Pohl CS, McGrath SD, Wendl MC, Zhang QY, Locke DP, et al: BreakDancer: an algorithm for high-resolution mapping of genomic structural variation. Nature Methods 2009, 6: 677-+.

16. Ye K, Schulz MH, Long Q, Apweiler R, Ning ZM: Pindel: a pattern growth approach to detect break points of large deletions and medium sized insertions from paired-end short reads. Bioinformatics 2009, 25: 2865–2871.

17. Chen K, Chen L, Fan X, Wallis J, Ding L, Weinstock G: TIGRA: A targeted iterative graph routing assembler for breakpoint assembly. Genome Research 2014, 24: 310–317.

18. Hormozdiari F, Hajirasouliha I, Dao P, Hach F, Yorukoglu D, Alkan C, Eichler EE, Sahinalp SC: Next-generation VariationHunter: combinatorial algorithms for transposon insertion discovery. Bioinformatics 2010, 26: i350–i357.

19. Jiang Y, Wang YD, Brudno M: PRISM: Pair-read informed split-read mapping for base-pair level detection of insertion, deletion and structural variants. Bioinformatics 2012, 28: 2576–2583.

20. Rausch T, Zichner T, Schlattl A, Stutz AM, Benes V, Korbel JO: DELLY: structural variant discovery by integrated paired-end and split-read analysis. Bioinformatics 2012, 28: I333–I339.

21. English AC, Salerno WJ, Hampton OA, Gonzaga-Jauregui C, Ambreth S, Ritter DI, Beck CR, Davis CF, Dahdouli M, Ma S, et al: Assessing structural variation in a personal genome-towards a human reference diploid genome. Bmc Genomics 2015, 16.

22. Tattini L, D’Aurizio R, Magi A: Detection of Genomic Structural Variants from Next-Generation Sequencing Data. Front Bioeng Biotechnol 2015, 3: 92.

23. Roberts RJ, Carneiro MO, Schatz MC: The advantages of SMRT sequencing. Genome Biol 2013, 14: 405.

24. Jain M, Olsen HE, Paten B, Akeson M: The Oxford Nanopore MinION: delivery of nanopore sequencing to the genomics community. Genome Biol 2016, 17: 239.

25. Sedlazeck FJ, Lee H, Darby CA, Schatz MC: Piercing the dark matter: bioinformatics of long-range sequencing and mapping. Nat Rev Genet 2018, 19: 329–346.

26. Goodwin S, McPherson JD, McCombie WR: Coming of age: ten years of next-generation sequencing technologies. Nat Rev Genet 2016, 17: 333–351.

27. English AC, Salerno WJ, Reid JG: PBHoney: identifying genomic variants via long-read discordance and interrupted mapping. BMC Bioinformatics 2014, 15: 180.

28. Huddleston J, Chaisson MJP, Steinberg KM, Warren W, Hoekzema K, Gordon D, Graves-Lindsay TA, Munson KM, Kronenberg ZN, Vives L, et al: Discovery and genotyping of structural variation from long-read haploid genome sequence data. Genome Research 2017, 27: 677–685.

29. Sedlazeck FJ, Rescheneder P, Smolka M, Fang H, Nattestad M, von Haeseler A, Schatz MC: Accurate detection of complex structural variations using single-molecule sequencing. Nature Methods 2018, 15: 461-+.

30. Heller D, Vingron M: SVIM: Structural Variant Identification using Mapped Long Reads. Bioinformatics 2019.

31. Ho SS, Urban AE, Mills RE: Structural variation in the sequencing era. Nat Rev Genet 2019.

32. Chaisson MJ, Tesler G: Mapping single molecule sequencing reads using basic local alignment with successive refinement (BLASR): application and theory. Bmc Bioinformatics 2012, 13.

33. Li H: Minimap2: pairwise alignment for nucleotide sequences. Bioinformatics 2018, 34: 3094–3100.

34. Jiang T, Liu B, Li J, Wang Y: rMETL: sensitive mobile element insertion detection with long read realignment. Bioinformatics 2019, 35: 3484–3486.

35. Jiang T, Fu YL, Liu B, Wang YD: Long-Read Based Novel Sequence Insertion Detection With rCANID. Ieee Transactions on Nanobioscience 2019, 18: 343–352.

36. Shao H, Ganesamoorthy D, Duarte T, Cao MD, Hoggart CJ, Coin LJM: npInv: accurate detection and genotyping of inversions using long read sub-alignment. BMC Bioinformatics 2017, 19: 261.

37. Zook JM, Catoe D, McDaniel J, Vang L, Spies N, Sidow A, Weng Z, Liu Y, Mason CE, Alexander N, et al: Extensive sequencing of seven human genomes to characterize benchmark reference materials. Sci Data 2016, 3: 160025.

38. Zook JM, Hansen NF, Olson ND, Chapman LM, Mullikin JC, Xiao C, Sherry S, Koren S, Phillippy AM, Boutros PC, et al: A robust benchmark for germline structural variant detection. bioRxiv 2019:664623.

39. Travers KJ, Chin CS, Rank DR, Eid JS, Turner SW: A flexible and efficient template format for circular consensus sequencing and SNP detection. Nucleic Acids Research 2010, 38.

40. Wenger AM, Peluso P, Rowell WJ, Chang PC, Hall RJ, Concepcion GT, Ebler J, Fungtammasan A, Kolesnikov A, Olson ND, et al: Accurate circular consensus long-read sequencing improves variant detection and assembly of a human genome. Nat Biotechnol 2019.

41. Clarke L, Fairley S, Zheng-Bradley X, Streeter I, Perry E, Lowy E, Tasse AM, Flicek P: The international Genome sample resource (IGSR): A worldwide collection of genome variation incorporating the 1000 Genomes Project data. Nucleic Acids Res 2017, 45: D854–D859.

42. Chaisson MJP, Sanders AD, Zhao X, Malhotra A, Porubsky D, Rausch T, Gardner EJ, Rodriguez OL, Guo L, Collins RL, et al: Multi-platform discovery of haplotype-resolved structural variation in human genomes. Nat Commun 2019, 10: 1784.

