## Supplementaly Notes for "Long-read-based Human Genomic Structural Variation Detection with cuteSV"

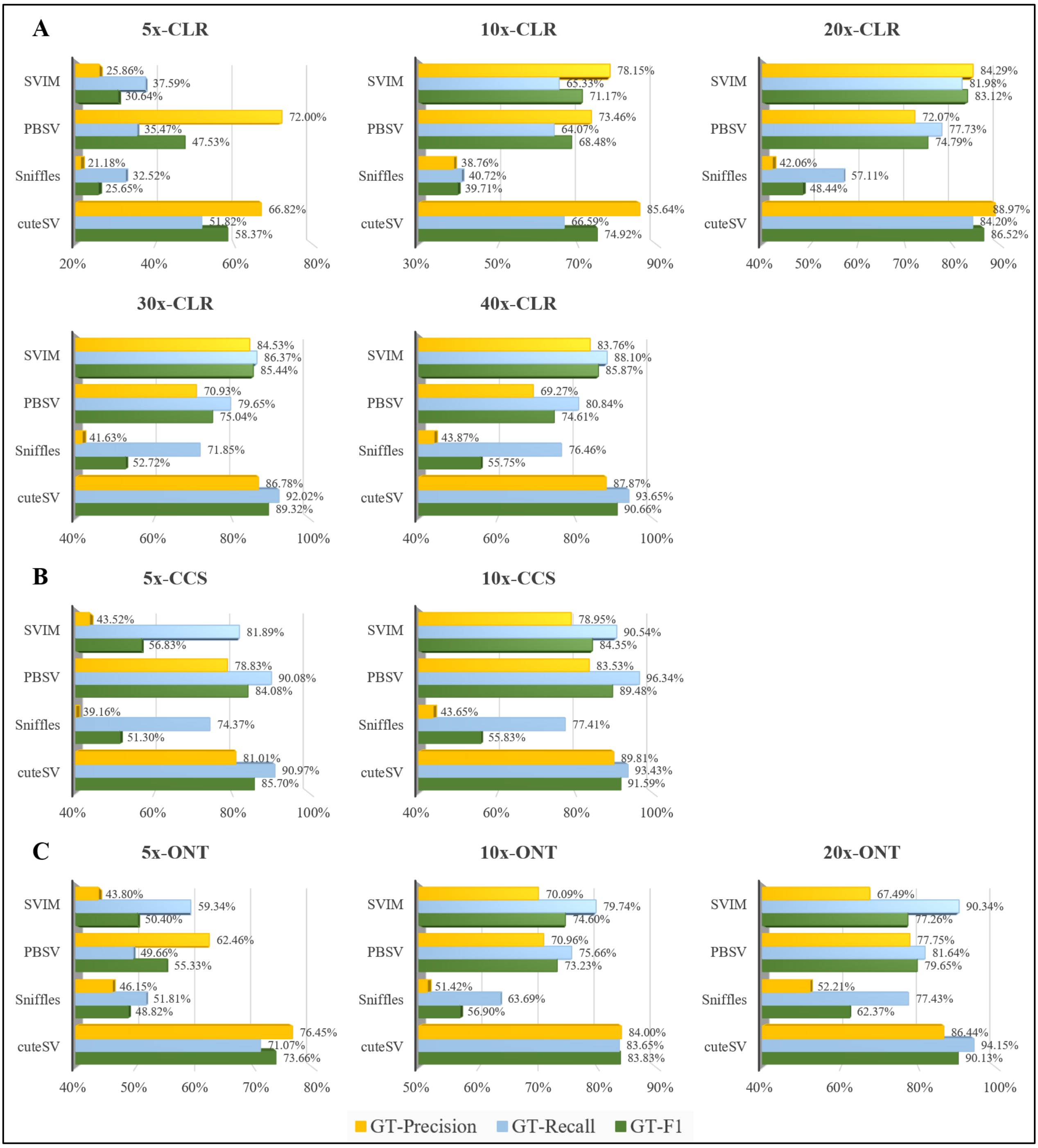

**Supplementary Figure 1. Benchmark results of the genotyping on HG002 down-sampling datasets.**

(**A**) GT-Precisions, GT-recalls and GT-F1 on HG002 PacBio CLR down-sampling datasets. (**B**) GT-Precisions, GT-recalls and GT-F1 on HG002 PacBio CCS down-sampling datasets. (**C**) GT-Precisions, GT-recalls and GT-F1 on HG002 ONT PromethION down-sampling datasets.

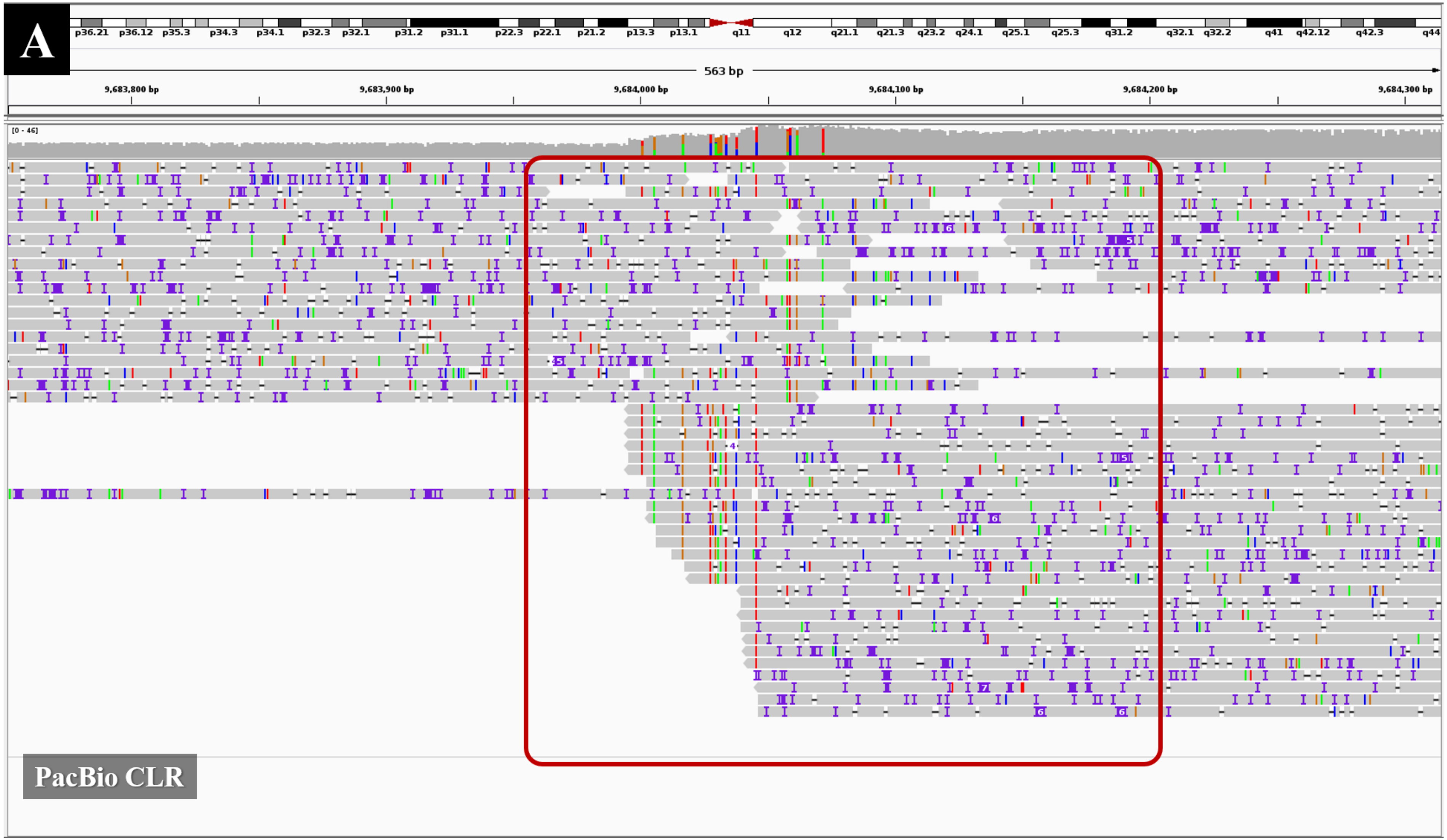
A
p36.21 p36.12 p35.3 p34.3 p34.1 p32.3 p32.1 p31.2 p31.1 p22.3 p22.1 p21.2 p13.3 p13.1 q11 q12 q21.1 q21.3 q23.2 q24.1 q25.1 q25.3 q31.2 q32.1 q32.2 q41 q42.12 q42.3 q44
563 bp
9,683,800 bp
9,683,900 bp
9,684,000 bp
9,684,100 bp
9,684,200 bp
9,684,300 bp
[0 - 46]
PacBio CLR

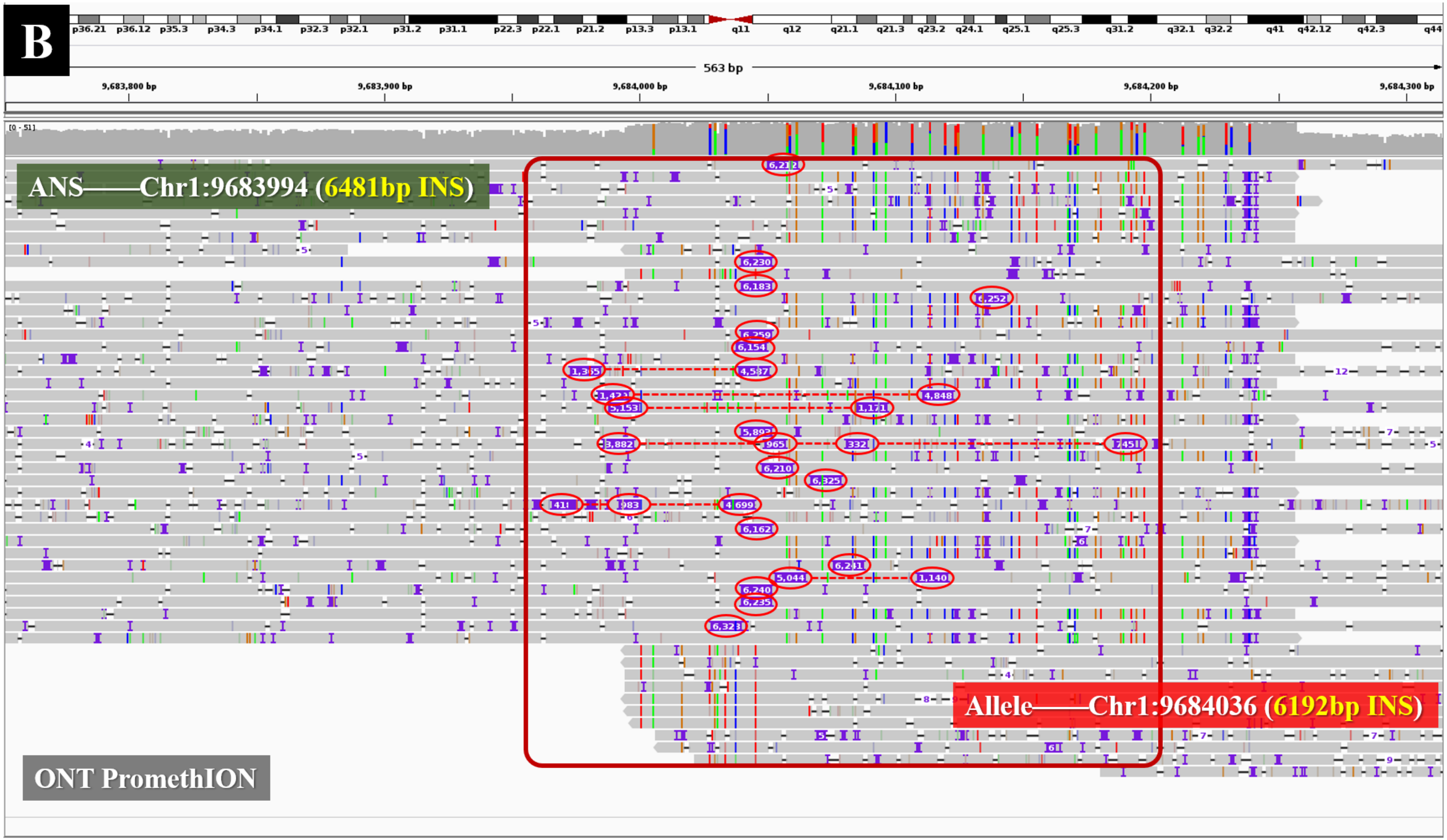

**Supplementary Figure 2. An example of an insertion only being detected with ONT PromethION data**

According to the ground truth, there is a 6481 bp insertion at chr1:9683994. (A) The Integrated Genomics Viewer (IGV) snapshot of the PacBio CLR read alignments at chr1:9683700-9684300 indicates that nearly 50% of the reads mapped to this region are about 5000 bp long and few of them are longer than 10 kbp. Due to the limited read length, it is difficult to produce alignments with such a large insertion, so none of the PacBio reads have an obvious insertion signature and cuteSV (as well as other SV callers) failed for this SV event. (B) The IGV snapshot of the ONT read alignments indicates that the minimum and average lengths of the reads mapped to this region are 18kbp and 58kbp, respectively. With superior read lengths, 20 of the reads were aligned with >6000 bp insertion in their CIGARs, which are very useful SV signatures. cuteSV captured these signatures successfully, and made a 6192 bp insertion at chr1:9684036, which highly coincides with the ground truth.

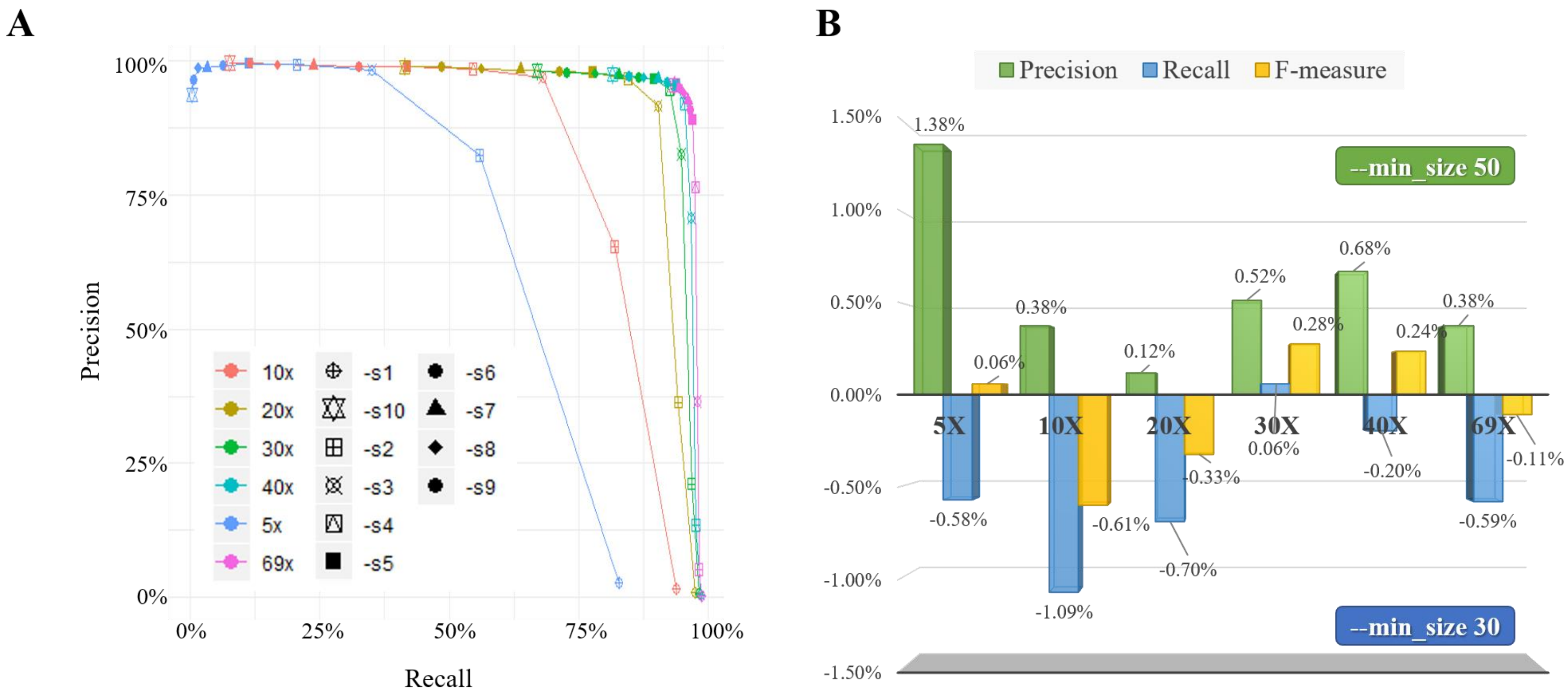

**Supplementary Figure 3. Results of cuteSV with various configurations on the parameters *--min_support* and *–min_size***

(A) The precision and recall for various sequencing coverages and the configurations of --*min_support* parameter. (B) The differences in precision, recall, and F-measure between two different configurations of –*min_size* parameter (–*min_size* = 30 and –*min_size* = 50). It is worth noting that the positive values indicate that cuteSV achieved higher statistics with –*min_size* = 50, and the negative values indicate higher statistics with –*min_size* = 30.

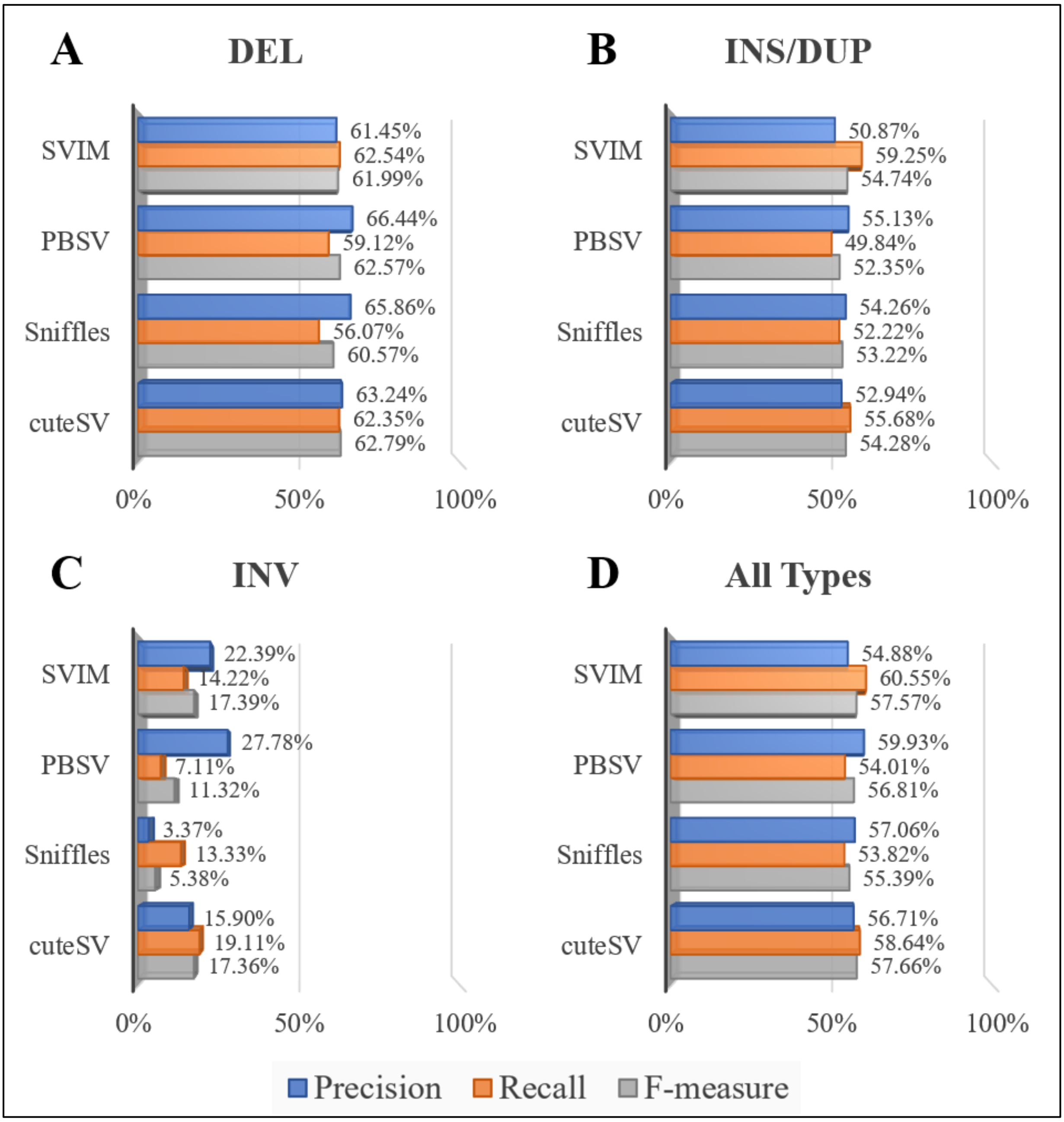

**Supplementary Figure 4. Results of the SV callers on the PacBio CLR dataset from NA19240 sample**

The subplots depict the precision, recalls and F-measures of the benchmarked SV callers for various types of SVs, i.e., (A) deletions, (B) insertions and duplications, (C) inversions and (D) all types of SVs.

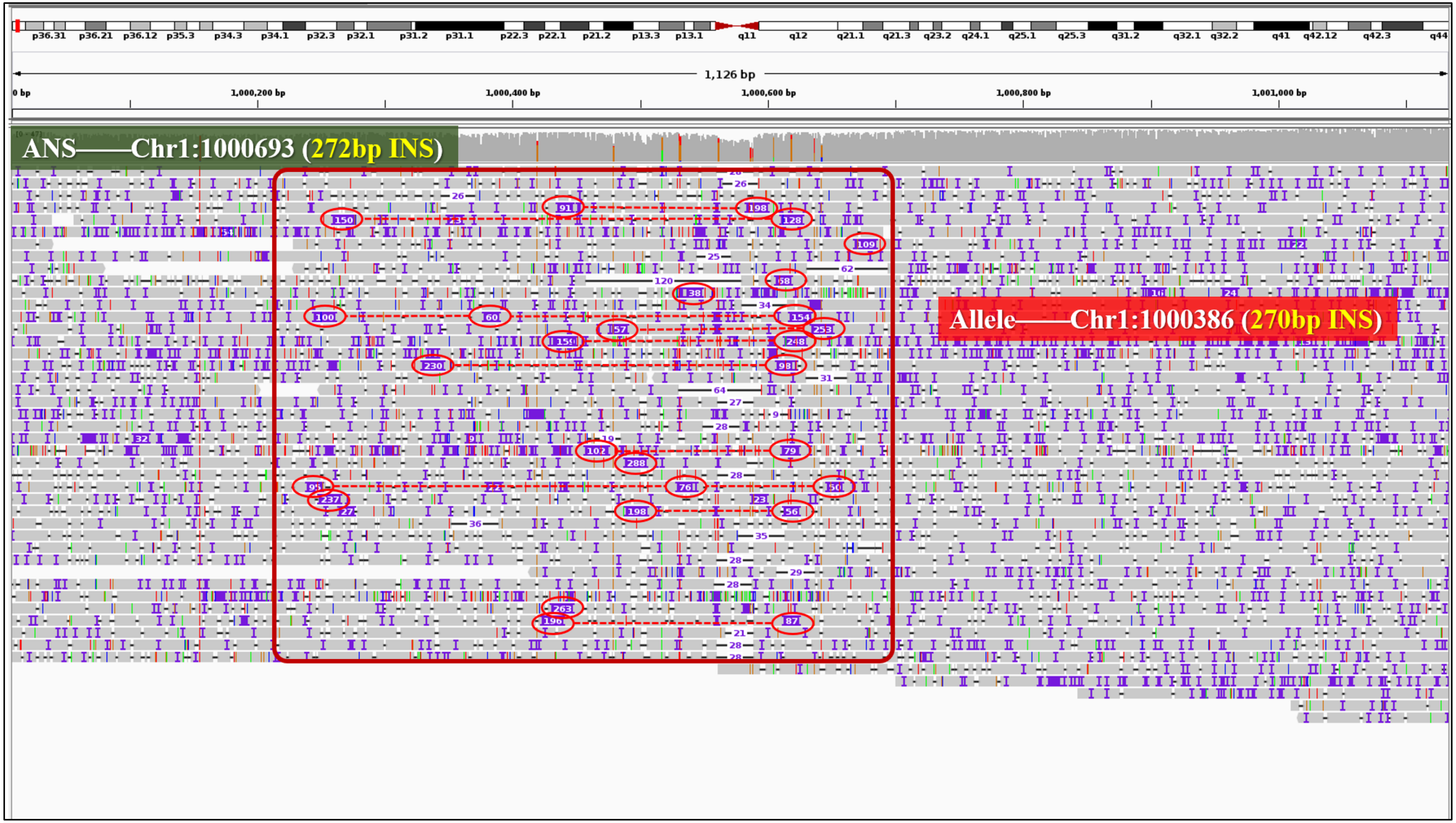

**Supplementary Figure 5. An example of fragile read alignments around insertion events**

According to the ground truth, there is a 272 bp insertion whose breakpoint is at chr1:1000693. The IGV snapshot of the PacBio CLR read alignments around this event is shown in the figure. It can be seen that many of the reads mapped to this region have insertion signatures. However, most of them have two nearby large insertions in their CIGARs instead of a large insertion (indicated by the red circles and dashed lines in the figure). In this case, cuteSV recognized 28 insertion signatures longer than 30 bp in length, and 22 of them satisfied the signature merging rule. cuteSV merged the split signatures to recover the signatures of the real larger insertion, which helps to make an insertion call (270 bp insertion at chr1:1000386) coinciding with the ground truth.

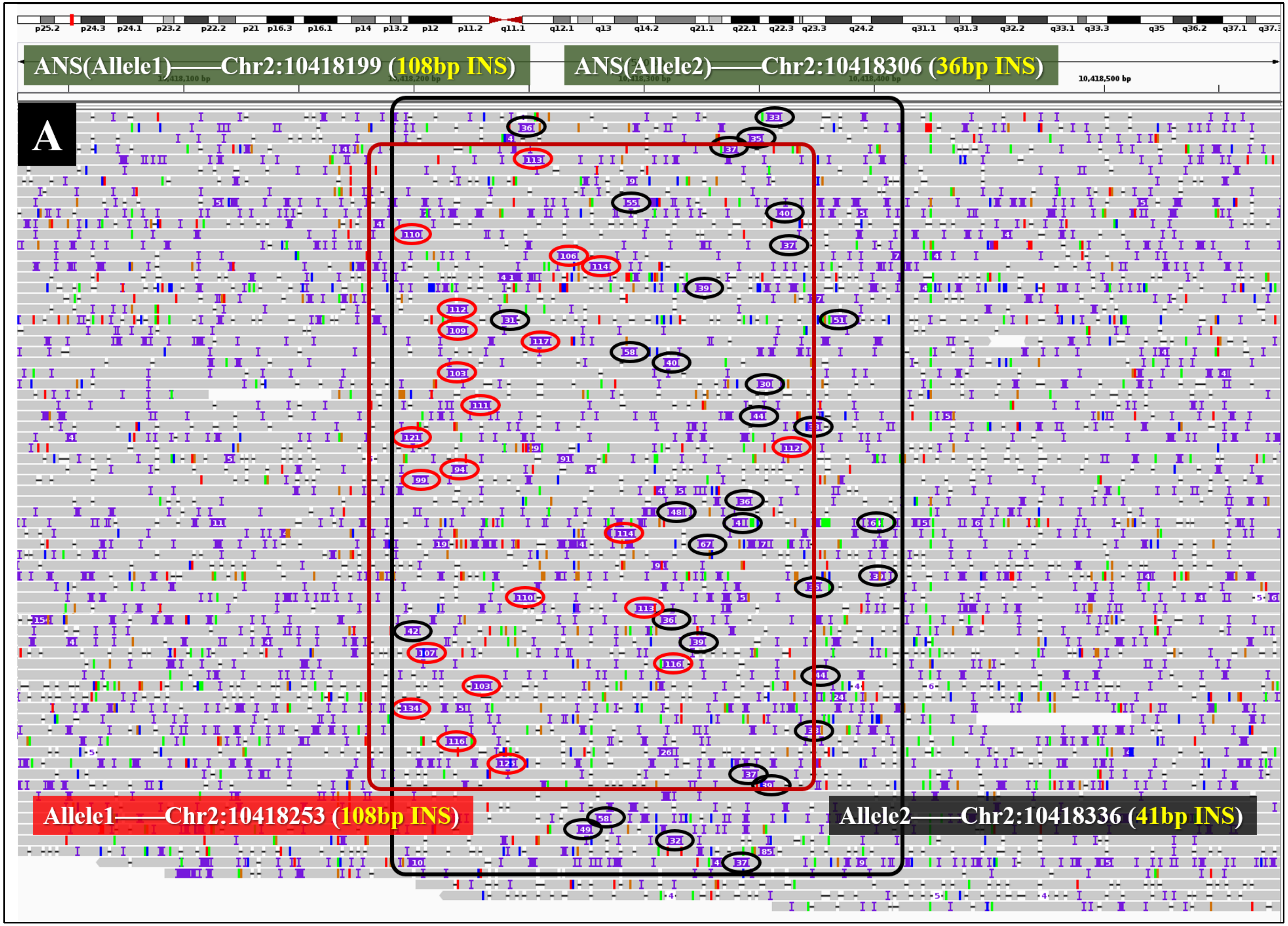
A
ANS(Allele1)——Chr2:10418199 (108bp INS)
ANS(Allele2)——Chr2:10418306 (36bp INS)
Allele1——Chr2:10418253 (108bp INS)
Allele2——Chr2:10418336 (41bp INS)

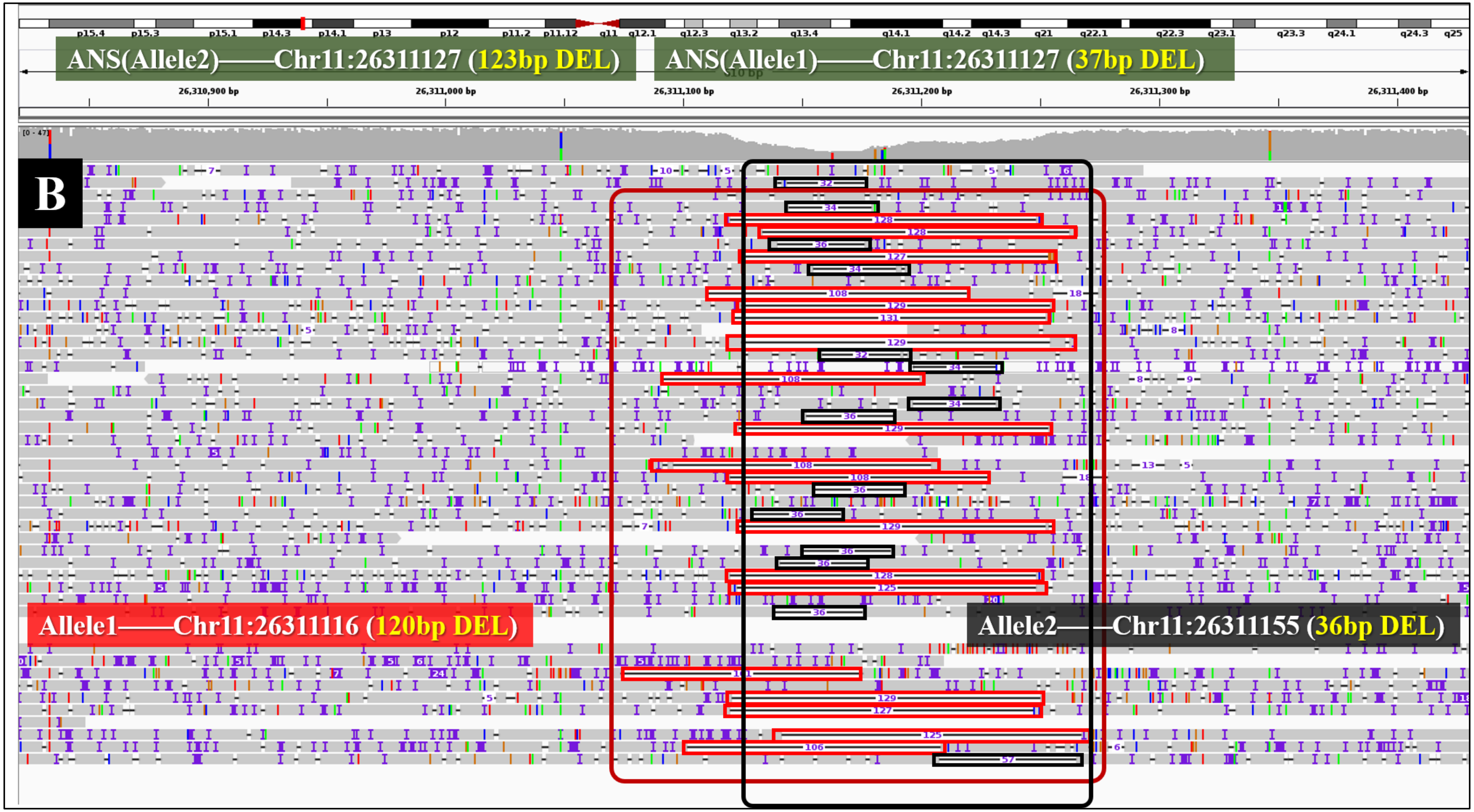

**Supplementary Figure 6. Two examples of the read alignments around multi-allelic SVs**

(A) According to the ground truth, there are two different insertions, i.e., a 108 bp insertion at chr2:10418199 and a 36 bp insertion at chr2:10418306. The IGV snapshot of the PacBio CLR read alignments around these events is shown in the figure. Many reads can be found mapped to this region with insertions of about 100 bp (marked with the red circle) and 40 bp (marked with the black circle) in their CIGARs. In this case, cuteSV called two different SVs, i.e., a 108 bp insertion at chr2:10418253 and a 41 bp insertion at chr2:10418336. (B) Similarly, there are two ground truth deletions, i.e., a 123 bp deletion at chr11:26311127 and a 37 bp deletion at chr11:26311127. cuteSV recognized one set of deletion signatures around 120 bp (marked with the red box) and one set of deletion signatures around 36 bp (marked with the black box). As a result, cuteSV made two deletion calls, i.e., a 120 bp at chr11:26311116 and a 36 bp at chr11:26311155. All these SV calls coincide with the ground truth.

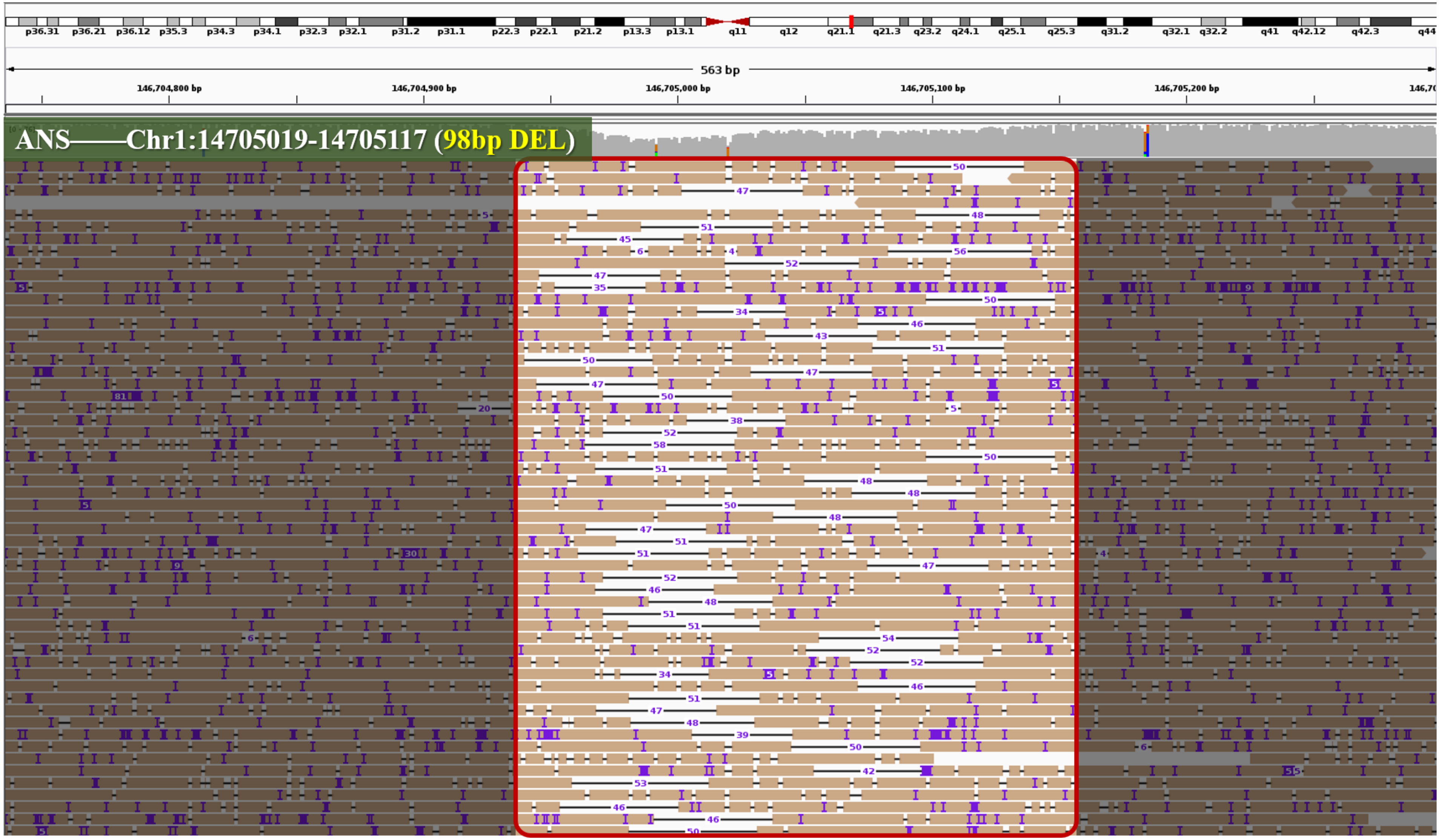

**Supplementary Figure 7. An example of a false-negative deletion**

According to the ground truth, there is a 98 bp deletion at chr1:14705019-14705117. The IGV snapshot of the PacBio CLR read alignments around this event is shown in the figure. In this region, there are only reads with 30 to 50 bp deletion in their CIGARs. Hence, cuteSV made a deletion call of about 50 bp. Meanwhile, none of the other benchmarked SV callers detected the 98bp deletion.

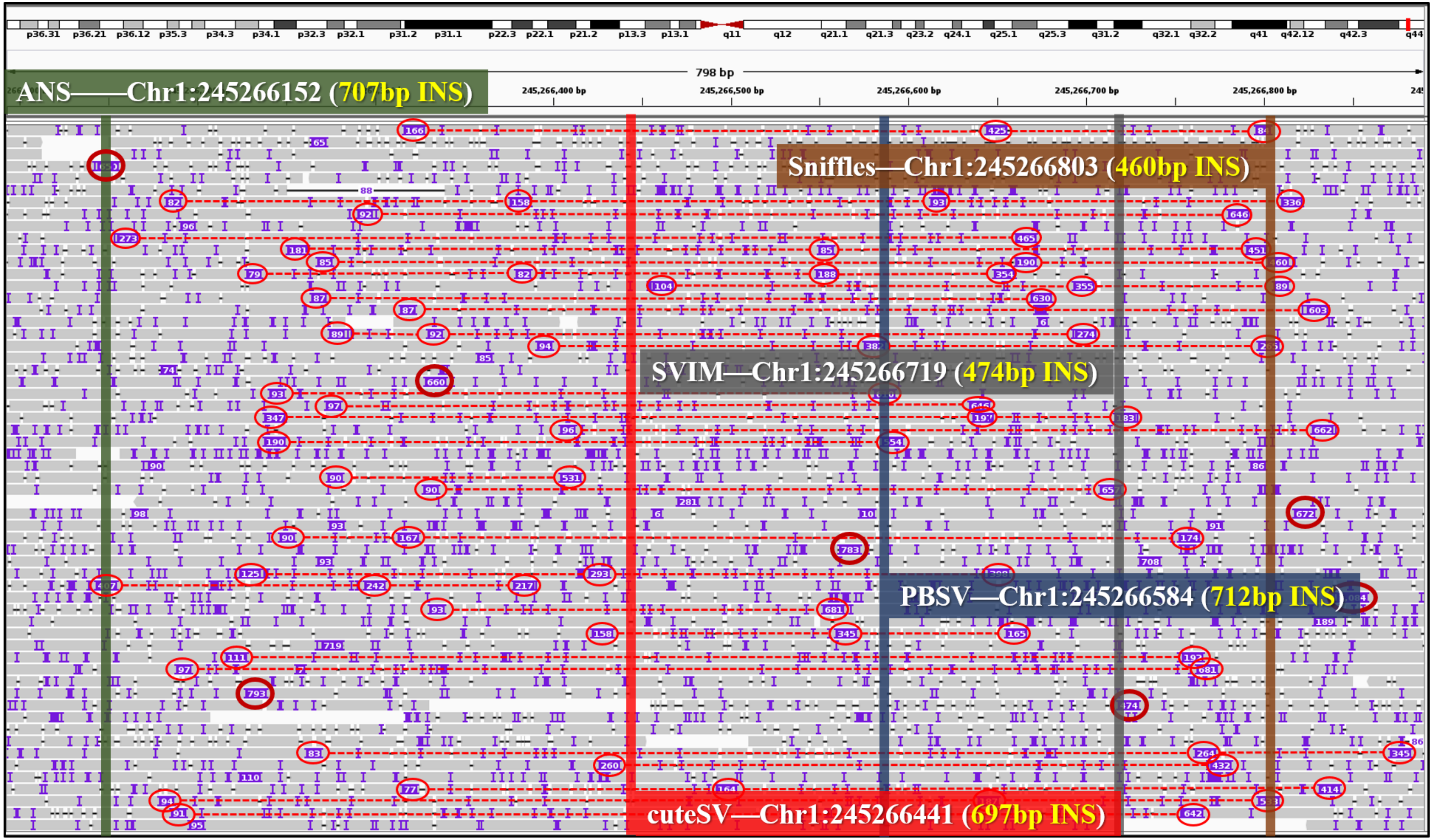

ANS——Chr1:245266152 (707bp INS)
Sniffles—Chr1:245266803 (460bp INS)
SVIM—Chr1:245266719 (474bp INS)
PBSV—Chr1:245266584 (712bp INS)
cuteSV—Chr1:245266441 (697bp INS)
798 bp
245,266,400 bp
245,266,500 bp
245,266,600 bp
245,266,700 bp
245,266,800 bp

**Supplementary Figure 8. An example of a false-negative insertion**

According to the ground truth, there is a 707 bp insertion at chr1:245266152. The IGV snapshot of the PacBio CLR read alignments around this event is shown in the figure. In this region, cuteSV recognized a lot of linked fragile insertion signatures (marked by red dashed lines in the figure) and used them to make a 697 bp insertion at chr1:245266441. However, the breakpoint of this call is 289 bp away from the breakpoint of a nearby ground truth call, so Truvari determined that the ground truth call is a false negative one. Moreover, PBSV made a 712 bp insertion at chr1:245266584, also away from the ground truth (by 432bp). For Sniffles and SVIM, their calls are even more different from the ground truth.

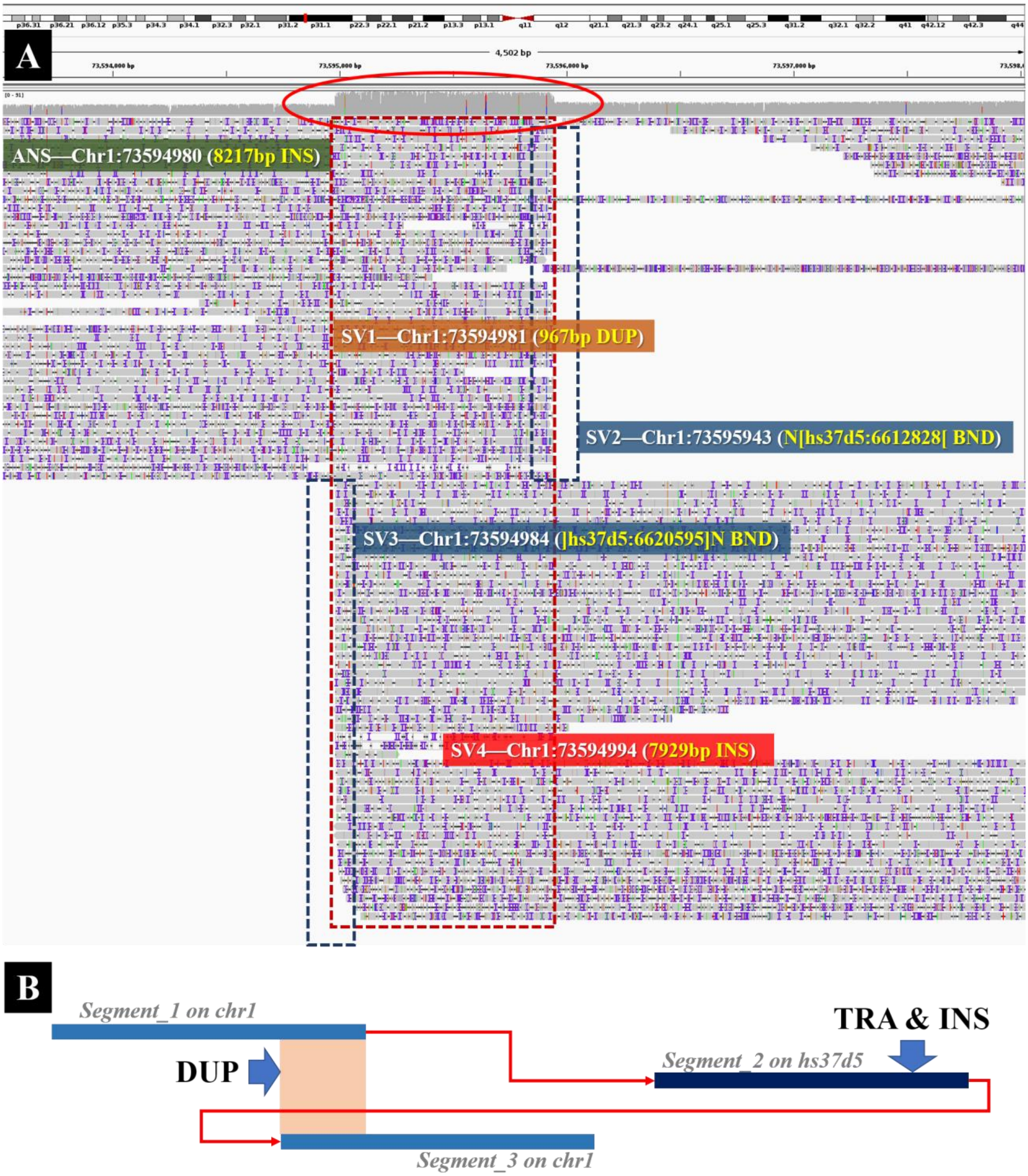

**Supplementary Figure 9. An example of the alignments of a "copy-and-paste" insertion with duplication**

(A) The snapshot of IGV for the alignments of ONT PromethION reads around a ground truth insertion (8217 bp insertion, breakpoint at chr1:73594980). In this local region, cuteSV recognized the signatures of translocation (marked by the deep blue dashed box) and made two BND calls, i.e., chr1:73595943(N[hs37d5:6612828) and chr1:73594984(]hs37d5:6620595]N). Moreover, cuteSV also recognized the signatures of duplication (marked by the red dotted box) and made a duplication call (duplication at chr1:73594981). (B) A schematic illustration of the complex SV event. This event can be seen as a local arrangement of three genomic segments (marked as segment_1, segment_2 and segment_3 in the figure). It is observed that segment_2 is a segment which can be aligned to a decoy sequence of the reference (hs37d5). So, this case can be considered as a copy-and-paste event, i.e., segment_2 is copied from the decoy sequence and pasted in between segment_1 and segment_3. In this situation, cuteSV skips the translocated segment (segment_2) and sees it as an insertion (7929 bp insertion at chr1:73594994) between segment_1 and segment_3.

**Supplementary Table 1. Benchmark results of different sequencing datasets on HG002**

| Dataset | Deletion | Insertion | Duplication | Inversion | Translocation | ALL |
|---|---|---|---|---|---|---|
| PacBio | 9634 | 14272 | 450 | 425 | 1824 | 26605 |
| ONT | 11005 | 13734 | 423 | 82 | 1183 | 26427 |
| High_conf | 4199 | 5442 | 0 | 0 | 0 | 9641 |
| PacBio ∩ ONT | 8088 | 10323 | 82 | 53 | 766 | 19312 |
| PacBio ∩ High_conf | 4044 | 5027 | 0 | 0 | 0 | 9071 |
| ONT ∩ High_conf | 4122 | 5192 | 0 | 0 | 0 | 9314 |
| PacBio ∩ ONT ∩ High_conf | 3993 | 4882 | 0 | 0 | 0 | 8875 |

PacBio and ONT indicate the SV callsets of cuteSV generated from the corresponding sequencing platforms. High_conf means the SV ground truth set of the HG002 human sample from GIAB. The assessment was performed using *cmp_hg2_platforms.py*, available at https://github.com/tjiangHIT/cuteSV.

**Supplementary Table 2. The number of false negative calls of HG002 sample under different methods**

| SV size | PBMM2 | | | | NGMLR | | | |
|---|---|---|---|---|---|---|---|---|
| | cuteSV | Sniffles | PBSV | SVIM | cuteSV | Sniffles | PBSV | SVIM |
| (-∞, -10000] | 1 | 12 | 1 | 2 | 2 | 0 | 3 | 3 |
| (-10000, -9000] | 1 | 4 | 1 | 2 | 1 | 0 | 0 | 1 |
| (-9000, -8000] | 0 | 2 | 1 | 0 | 1 | 0 | 0 | 2 |
| (-8000, -7000] | 0 | 0 | 0 | 0 | 0 | 0 | 0 | 0 |
| (-7000, -6000] | 1 | 32 | 2 | 6 | 1 | 1 | 3 | 2 |
| (-6000, -5000] | 1 | 23 | 0 | 7 | 3 | 3 | 1 | 4 |
| (-5000, -4000] | 3 | 24 | 2 | 10 | 2 | 1 | 2 | 4 |
| (-4000, -3000] | 11 | 45 | 4 | 24 | 0 | 0 | 1 | 7 |
| (-3000, -2000] | 18 | 51 | 8 | 34 | 4 | 4 | 3 | 6 |
| (-2000, -1000] | 57 | 78 | 44 | 72 | 4 | 5 | 11 | 6 |
| (-1000, -900] | 4 | 5 | 5 | 4 | 1 | 2 | 4 | 2 |
| (-900, -800] | 1 | 2 | 6 | 1 | 0 | 1 | 2 | 2 |
| (-800, -700] | 5 | 4 | 10 | 7 | 3 | 0 | 4 | 1 |
| (-700, -600] | 4 | 5 | 8 | 4 | 0 | 2 | 2 | 1 |
| (-600, -500] | 3 | 4 | 6 | 2 | 2 | 2 | 2 | 2 |
| (-500, -400] | 0 | 2 | 9 | 5 | 0 | 2 | 2 | 3 |
| (-400, -300] | 5 | 8 | 32 | 7 | 6 | 14 | 28 | 7 |
| (-300, -200] | 6 | 6 | 20 | 5 | 5 | 5 | 10 | 5 |
| (-200, -100] | 9 | 19 | 29 | 8 | 9 | 28 | 43 | 11 |
| (-100, -50] | 25 | 185 | 93 | 30 | 38 | 260 | 101 | 38 |
| [50, 100) | 24 | 90 | 164 | 24 | 29 | 158 | 140 | 38 |
| [100, 200) | 16 | 36 | 85 | 27 | 20 | 45 | 90 | 26 |
| [200, 300) | 8 | 24 | 42 | 18 | 23 | 36 | 140 | 30 |
| [300, 400) | 6 | 21 | 104 | 20 | 43 | 46 | 375 | 46 |
| [400, 500) | 11 | 18 | 30 | 20 | 30 | 21 | 53 | 20 |
| [500, 600) | 8 | 19 | 14 | 12 | 22 | 17 | 38 | 16 |
| [600, 700) | 7 | 16 | 21 | 14 | 30 | 19 | 41 | 16 |
| [700, 800) | 13 | 13 | 17 | 15 | 15 | 17 | 25 | 20 |
| [800, 900) | 7 | 13 | 22 | 18 | 13 | 13 | 22 | 13 |
| [900, 1000) | 6 | 13 | 13 | 14 | 13 | 15 | 12 | 13 |
| [1000, 2000) | 110 | 169 | 154 | 204 | 83 | 181 | 89 | 207 |
| [2000, 3000) | 49 | 130 | 46 | 143 | 35 | 110 | 19 | 150 |
| [3000, 4000) | 35 | 80 | 28 | 78 | 29 | 68 | 20 | 78 |
| [4000, 5000) | 26 | 47 | 12 | 46 | 26 | 44 | 15 | 45 |
| [5000, 6000) | 9 | 18 | 6 | 18 | 9 | 17 | 9 | 18 |
| [6000, 7000) | 40 | 58 | 36 | 58 | 47 | 55 | 34 | 58 |
| [7000, 8000) | 5 | 14 | 6 | 14 | 11 | 15 | 13 | 14 |
| [8000, 9000) | 7 | 7 | 6 | 7 | 7 | 7 | 7 | 7 |
| [9000, 10000) | 5 | 5 | 5 | 5 | 5 | 5 | 5 | 5 |
| [10000, +∞) | 23 | 22 | 24 | 22 | 23 | 22 | 24 | 22 |

**Supplementary Table 3. The number of false positive calls of HG002 sample under different methods**

| SV size | PBMM2 | | | | NGMLR | | | |
|---|---|---|---|---|---|---|---|---|
| | cuteSV | Sniffles | PBSV | SVIM | cuteSV | Sniffles | PBSV | SVIM |
| (-∞, -10000] | 2 | 1 | 7 | 1 | 3 | 1 | 3 | 1 |
| (-10000, -9000] | 0 | 0 | 0 | 0 | 1 | 0 | 0 | 1 |
| (-9000, -8000] | 2 | 0 | 2 | 2 | 2 | 1 | 1 | 1 |
| (-8000, -7000] | 0 | 0 | 1 | 0 | 0 | 0 | 0 | 0 |
| (-7000, -6000] | 1 | 0 | 2 | 1 | 1 | 0 | 0 | 1 |
| (-6000, -5000] | 0 | 0 | 2 | 1 | 2 | 1 | 1 | 2 |
| (-5000, -4000] | 0 | 0 | 1 | 0 | 0 | 0 | 0 | 0 |
| (-4000, -3000] | 1 | 0 | 3 | 1 | 0 | 0 | 0 | 0 |
| (-3000, -2000] | 2 | 0 | 2 | 2 | 0 | 1 | 2 | 0 |
| (-2000, -1000] | 0 | 1 | 5 | 0 | 1 | 5 | 12 | 4 |
| (-1000, -900] | 1 | 1 | 1 | 1 | 0 | 0 | 1 | 0 |
| (-900, -800] | 0 | 2 | 4 | 0 | 1 | 0 | 5 | 1 |
| (-800, -700] | 1 | 0 | 2 | 0 | 0 | 0 | 0 | 0 |
| (-700, -600] | 1 | 2 | 5 | 3 | 0 | 0 | 3 | 3 |
| (-600, -500] | 0 | 1 | 5 | 1 | 0 | 0 | 3 | 2 |
| (-500, -400] | 1 | 3 | 4 | 3 | 1 | 1 | 4 | 5 |
| (-400, -300] | 4 | 4 | 16 | 2 | 0 | 0 | 18 | 3 |
| (-300, -200] | 4 | 11 | 13 | 11 | 5 | 4 | 14 | 4 |
| (-200, -100] | 15 | 25 | 41 | 30 | 9 | 18 | 22 | 19 |
| (-100, -50] | 69 | 59 | 97 | 77 | 47 | 46 | 77 | 60 |
| [50, 100) | 273 | 187 | 282 | 286 | 205 | 151 | 250 | 215 |
| [100, 200) | 67 | 120 | 51 | 121 | 48 | 74 | 40 | 109 |
| [200, 300) | 23 | 58 | 12 | 48 | 6 | 44 | 5 | 89 |
| [300, 400) | 17 | 33 | 12 | 34 | 9 | 64 | 2 | 57 |
| [400, 500) | 3 | 26 | 5 | 18 | 0 | 26 | 6 | 48 |
| [500, 600) | 4 | 10 | 10 | 11 | 1 | 24 | 27 | 16 |
| [600, 700) | 5 | 4 | 5 | 8 | 1 | 15 | 14 | 19 |
| [700, 800) | 0 | 3 | 4 | 4 | 0 | 17 | 19 | 12 |
| [800, 900) | 0 | 2 | 1 | 2 | 0 | 8 | 12 | 10 |
| [900, 1000) | 0 | 4 | 3 | 0 | 0 | 9 | 8 | 2 |
| [1000, 2000) | 3 | 2 | 17 | 6 | 7 | 22 | 27 | 13 |
| [2000, 3000) | 1 | 1 | 6 | 3 | 1 | 2 | 5 | 1 |
| [3000, 4000) | 0 | 0 | 5 | 0 | 0 | 3 | 1 | 0 |
| [4000, 5000) | 0 | 0 | 2 | 0 | 0 | 0 | 0 | 0 |
| [5000, 6000) | 0 | 1 | 0 | 2 | 0 | 1 | 0 | 1 |
| [6000, 7000) | 0 | 0 | 0 | 1 | 0 | 1 | 0 | 0 |
| [7000, 8000) | 0 | 0 | 0 | 0 | 0 | 0 | 0 | 0 |
| [8000, 9000) | 0 | 0 | 0 | 0 | 0 | 0 | 0 | 0 |
| [9000, 10000) | 0 | 0 | 0 | 0 | 0 | 0 | 0 | 0 |
| [10000, +∞) | 0 | 0 | 0 | 0 | 0 | 0 | 0 | 0 |

**Supplementary Table 4. The precision and recall of HG002 sample for various sequencing coverages and the configurations of *--min_support* parameter**

| Coverage | Support-read | Precision | Recall | Coverage | Support-read | Precision | Recall |
|---|---|---|---|---|---|---|---|
| 5× | -s1 | 2.38% | 83.27% | 30× | -s1 | 0.50% | 97.79% |
| | -s2 | 80.50% | 56.45% | | -s2 | 19.13% | 96.26% |
| | -s3 | 98.27% | 35.95% | | -s3 | 77.87% | 94.48% |
| | -s4 | 99.27% | 21.16% | | -s4 | 93.50% | 92.55% |
| | -s5 | 99.30% | 11.75% | | -s5 | 96.10% | 89.92% |
| | -s6 | 99.08% | 6.69% | | -s6 | 96.36% | 87.53% |
| | -s7 | 98.53% | 3.47% | | -s7 | 96.83% | 84.04% |
| | -s8 | 99.39% | 1.70% | | -s8 | 97.17% | 79.65% |
| | -s9 | 98.41% | 0.64% | | -s9 | 97.50% | 74.53% |
| | -s10 | 96.97% | 0.32% | | -s10 | 97.66% | 69.17% |
| 10× | -s1 | 1.37% | 93.72% | 40× | -s1 | 0.39% | 98.28% |
| | -s2 | 62.30% | 82.48% | | -s2 | 12.19% | 97.30% |
| | -s3 | 96.19% | 69.12% | | -s3 | 63.49% | 96.55% |
| | -s4 | 98.13% | 55.66% | | -s4 | 88.83% | 95.37% |
| | -s5 | 98.62% | 42.91% | | -s5 | 94.18% | 94.05% |
| | -s6 | 98.75% | 33.72% | | -s6 | 94.90% | 92.70% |
| | -s7 | 98.81% | 25.06% | | -s7 | 95.79% | 91.12% |
| | -s8 | 99.19% | 17.81% | | -s8 | 96.40% | 88.89% |
| | -s9 | 99.25% | 12.42% | | -s9 | 96.72% | 86.24% |
| | -s10 | 99.38% | 8.34% | | -s10 | 96.88% | 83.22% |
| 20× | -s1 | 0.72% | 96.99% | 69× | -s1 | 0.25% | 98.54% |
| | -s2 | 33.46% | 94.05% | | -s2 | 4.96% | 97.93% |
| | -s3 | 89.70% | 90.48% | | -s3 | 28.47% | 97.62% |
| | -s4 | 96.21% | 85.21% | | -s4 | 63.80% | 97.33% |
| | -s5 | 97.36% | 78.83% | | -s5 | 82.12% | 96.88% |
| | -s6 | 97.63% | 72.92% | | -s6 | 83.79% | 96.51% |
| | -s7 | 97.85% | 65.69% | | -s7 | 88.21% | 96.18% |
| | -s8 | 98.03% | 58.21% | | -s8 | 91.53% | 95.57% |
| | -s9 | 98.33% | 50.69% | | -s9 | 93.79% | 94.91% |
| | -s10 | 98.43% | 43.49% | | -s10 | 94.78% | 94.09% |

**Supplementary Table 5. The summarization of precision, recall and F-measure on the 40× NA19240 PacBio CLR dataset**

| SV type | Tool | Precision | Recall | F-measure | Total calls | TP calls |
|---|---|---|---|---|---|---|
| Deletion | cuteSV | 63.24% | 62.35% | 62.79% | 11220 | 7096 |
| | Sniffles | 65.86% | 56.07% | 60.57% | 9787 | 6446 |
| | PBSV | 66.44% | 59.12% | 62.57% | 10367 | 6888 |
| | SVIM | 61.45% | 62.54% | 61.99% | 12113 | 7444 |
| Insertion & Duplication | cuteSV | 52.94% | 55.68% | 54.28% | 15728 | 8327 |
| | Sniffles | 54.26% | 52.22% | 53.22% | 14310 | 7765 |
| | PBSV | 55.13% | 49.84% | 52.35% | 13711 | 7559 |
| | SVIM | 50.87% | 59.25% | 54.74% | 18806 | 9567 |
| Inversion | cuteSV | 15.90% | 19.11% | 17.36% | 346 | 55 |
| | Sniffles | 3.37% | 13.33% | 5.38% | 860 | 29 |
| | PBSV | 27.78% | 7.11% | 11.32% | 54 | 15 |
| | SVIM | 22.39% | 14.22% | 17.39% | 134 | 30 |
| All types | cuteSV | 56.71% | 58.64% | 57.66% | 27294 | 15478 |
| | Sniffles | 57.06% | 53.82% | 55.39% | 24957 | 14240 |
| | PBSV | 59.93% | 54.01% | 56.81% | 24132 | 14462 |
| | SVIM | 54.88% | 60.55% | 57.57% | 31053 | 17041 |

The NA19240 raw PacBio CLR sequencing data was aligned with PBMM2. The evaluation was performed using *cmp_NA19240.py* available at https://github.com/tjiangHIT/cuteSV.

**Supplementary Table 6. The availability of raw sequencing datasets, alignments and truth sets**

| ID | Technology | Description | Availability |
|---|---|---|---|
| hs37d5 | NULL | Reference genome | ftp://ftp.1000genomes.ebi.ac.uk/vol1/ftp/technical/reference/phase2_reference_assembly_sequence |
| HG002 | PacBio CLR | Alignment | https://ftp.ncbi.nlm.nih.gov/giab/ftp/data/AshkenazimTrio/HG002_NA24385_son/PacBio_MtSinai_NIST/Baylor_NGMLR_bam_GRCh37/all_reads.fa.giab_h002_ngmlr-0.2.3_mapped.bam |
| HG003 | PacBio CLR | Alignment | https://ftp.ncbi.nlm.nih.gov/giab/ftp/data/AshkenazimTrio/HG003_NA24149_father/PacBio_MtSinai_NIST/Baylor_NGMLR_bam_GRCh37/all_reads.fa.giab_h003_ngmlr-0.2.3_mapped.bam |
| HG004 | PacBio CLR | Alignment | https://ftp.ncbi.nlm.nih.gov/giab/ftp/data/AshkenazimTrio/HG004_NA24143_mother/PacBio_MtSinai_NIST/Baylor_NGMLR_bam_GRCh37/all_reads.fa.giab_h004_ngmlr-0.2.3_mapped.bam |
| NA19240 | PacBio CLR | Alignment | ftp://ftp.1000genomes.ebi.ac.uk/vol1/ftp/data_collections/hgsv_sv_discovery/working/20160905_smithm_pacbio_aligns/NA19240_bwamem_GRCh38DH_YRI_20160905_pacbio.bam |
| HG002 | PacBio HiFi | Raw reads | ftp://ftp-trace.ncbi.nlm.nih.gov/giab/ftp/data/AshkenazimTrio/HG002_NA24385_son/PacBio_CCS_15kb/ |
| HG002 | Oxford Nanopore | Raw reads | ftp://ftp.ncbi.nlm.nih.gov/giab/ftp/data/AshkenazimTrio/HG002_NA24385_son/UCSC_Ultralong_OxfordNanopore_Promethion/ |
| HG002 | NULL | High confidence callsets | https://ftp-trace.ncbi.nlm.nih.gov/giab/ftp/data/AshkenazimTrio/analysis/NIST_SVs_Integration_v0.6/HG002_SVs_Tier1_v0.6.vcf.gz |
| HG002 | NULL | High confidence regions | https://ftp-trace.ncbi.nlm.nih.gov/giab/ftp/data/AshkenazimTrio/analysis/NIST_SVs_Integration_v0.6/HG002_SVs_Tier1_v0.6.bed |
| NA19240 | NULL | Callsets | ftp://ftp.ncbi.nlm.nih.gov/pub/dbVar/data/Homo_sapiens/by_study/vcf/nstd152.GRCh37.variant_call.vcf.gz |

### Supplementary Notes

#### 1. Commands used for read alignment and SV calling

##### 1.1 Extraction of raw sequencing long reads

samtools fasta {in.bam} > {raw_long_reads.fa}

##### 1.2 Read alignment

###### PBMM2 alignment

**Index**

pbmm2 index {reference.fa} {reference.mmi} (optional: --preset CCS)

**Align**

pbmm2 align {reference.mmi} {raw_long_reads.fa} {raw_long_reads_pbmm2.bam} --sort --rg '@RG\tID:${SAMPLE}' --sample {SAMPLE} (optional: --preset CCS)

###### NGMLR alignment

ngmlr -r {reference} -q {raw_long_reads.fa} -o {raw_long_reads_ngmlr.sam} (optional: -x ont)

###### Minimap2 alignment

minimap2 {reference} {raw_long_reads.fq} -a -z 600,200 -x map-ont –MD -Y -o {raw_long_reads_minimap2.sam} -R '@RG\tID:${SAMPLE}'

##### 1.3 Post-processing of read alignment

###### Alignment sorting

samtools view -buS {raw_long_reads_ngmlr.sam} | samtools sort -O bam -T {tempdir} - > {raw_long_reads_ngmlr.bam} && samtools index {raw_long_reads_ngmlr.bam}

###### Alignment down-sampling

samtools view -bS -s {ratio} {raw_long_reads.bam} > {raw_long_reads_subset.bam} && samtools index {raw_long_reads_subset.bam}

##### 1.4 SV calling

###### cuteSV calling

**For PacBio CLR and ONT data**

python3 cuteSV.py {raw_long_reads.bam} {raw_long_reads_cuteSV.vcf} {workdir} -s {min_support} -l {min_svsize} –max_cluster_bias_INS 100 –diff_ratio_merging_INS 0.2 –diff_ratio_filtering_INS 0.6 –diff_ratio_filtering_DEL 0.7

**For PacBio HiFi data**

python3 cuteSV.py {raw_long_reads.bam} {raw_long_reads_cuteSV.vcf} {workdir} -s {min_support} -l {min_svsize} –max_cluster_bias_INS 200 –diff_ratio_merging_INS 0.65 –diff_ratio_filtering_INS 0.65 –diff_ratio_filtering_DEL 0.35

###### Sniffles calling

sniffles -s {min_support} -l {min_svsize} -m {raw_long_reads.bam} -v {raw_long_reads_sniffles.vcf} (optional: --skip_parameter_estimation)

###### PBSV calling

pbsv discover {raw_long_reads.bam} {raw_long_reads.svsig.gz}

pbsv call {reference.fa} {raw_long_reads.svsig.gz} {raw_long_reads_pbsv.vcf} (optional: --ccs, -t INS,DEL)

##### SVIM calling

```
svim alignment –min_sv_size 30 {workdir} {raw_long_reads.bam} {reference}
cat {workdir/final_results.vcf} | awk '{ if($1 ~ /^#/) { print $0 } else { if($5=="<DEL>" || $5=="<INS>") { print $0 }}}' | grep -v
'SUPPORT=1;\|SUPPORT=2;\|SUPPORT=3;\|SUPPORT=4;\|SUPPORT=5;\|SUPPORT=6;\|SUPPORT=7;\|SUPPORT=8;\|SUPP
ORT=9;' | sed 's/DUP:INT/INS/g' | sed 's/DUP:TANDEM/INS/g' > {raw_long_reads_svim.vcf}
```

or

```
cat {workdir/final_results.vcf} | awk '{ if($1 ~ /^#/) { print $0 } else { if($5=="<DEL>" || $5=="<INS>") { print $0 }}}' | grep -v
'SUPPORT=1;\|SUPPORT=2;\|SUPPORT=3;\|SUPPORT=4;' | sed 's/DUP:INT/INS/g' | sed 's/DUP:TANDEM/INS/g' >
{raw_long_reads_svim.vcf}
```

##### 2. Truvari benchmarking

```
bgzip {raw_long_reads.vcf} && tabix {raw_long_reads.vcf.gz}
truvari -f {reference.fa} -b {HG002_SVs_Tier1_v0.6.vcf.gz} --includebed {HG002_SVs_Tier1_v0.6.bed} -o {benchmarkdir} –
passonly --giabreport -r 1000 -p 0.00 -c {raw_long_reads.vcf.gz}
```
